# Physiology mediates transcontinental radiations of freshwater fishes

**DOI:** 10.64898/2026.09.04.749521

**Authors:** Chase D. Brownstein, Richard C. Harrington, Thomas J. Near

## Abstract

Adaptive radiations are clades that have rapidly accumulated exceptional species richness and ecological diversity. The ability to access sources of ecological opportunity, such as new habitats, is thought to be essential for their success. Transcontinental lineages of exclusively freshwater fishes, such as the 1,700 species of cichlids distributed across Central and South America, Africa, India, and Madagascar, are some of the most celebrated adaptive radiations. Yet, what features facilitated their dispersal across marine barriers and diversification in geologically young freshwater ecosystems remain debated. Here, we show that a common mechanism, saltwater tolerance, provides an explanation for how species-rich transcontinental freshwater fish radiations, including cichlids, killifishes, pupfishes, and livebearers, accessed ecological opportunity and subsequently diversified. Marine salinity tolerance was retained after cichlids, killifishes and their relatives emerged from a rapid radiation during the Cretaceous-Paleogene boundary. The retention of this ancestral physiological trait throughout the last 45 million years of Earth history repeatedly enabled cichlids and other fishes to disperse across marine barriers and radiate in freshwater ecosystems on multiple continents, including geologically young features in Africa and North America that have repeatedly experienced saline conditions over tens of thousands of years of global climate change. These results show how the larger evolutionary context of adaptive radiations can sculpt their responses to ecological opportunity.

## Introduction

The ability to invade new habitats and ecological niches is an essential aspect of organismal biology that governs the generation of species diversity (*1–6*). Dispersals into new ecological contexts are implicated in shaping nearly all classic examples of adaptive radiations: the rapid accumulation of ecologically diverse species from a single common ancestor (*3*, *7*). Hawaiian Honeycreepers, Malagasy Vangas, and Darwin’s Finches (*8–10*), Andean lupins (*11*, *12*), and Antarctic notothenioid fishes (*5*, *13–15*) have all diversified in response to the opportunities afforded by geographic regions and habitats colonized by the common ancestors of these adaptive radiations (*1*, *7*).

Among adaptive radiations, few rival the species diversity and morphological disparity of fishes in the clade *Cichlidae* from Central and South America and the rift lakes of Africa (*16*, *17*), which comprise nearly 2,000 valid species (*18*) and many more awaiting formal description (*19*). Cichlids are a primary model system for the study of adaptive radiation, and the substantial fossil record (*6*, *20–24*) and extensive genomic resources (*17*, *25–36*) available for this clade has enabled detailed investigation of the factors underlying its species richness, ecological and behavioral diversity, and morphological disparity. The tempo and mode of cichlid adaptive radiation has been extensively studied, but in a manner largely independent of the second major historical controversy surrounding this clade: its age (*27*, *32*, *37–45*). The restriction of living cichlids to freshwater and brackish environments was classically interpreted as evidence that the transcontinental distribution of *Cichlidae* across the southern hemisphere reflects an ancient set of divergences that were driven by the Mesozoic fragmentation of the southern continents (*32*, *43*). This ancient vicariance hypothesis has been extended to other freshwater fish clades distributed across isolated landmasses (*39*, *41*, *43*, *46*), but has been undermined by the young ages recovered for the common ancestors of transcontinental fish clades in molecular phylogenies (*17*, *38*, *40*, *47–50*).

Yet, for cichlids and other transcontinental clades with exclusively freshwater distributions across their living and extinct diversity, it remains unclear how these dispersals took place. Classic studies of freshwater fish biogeography that predate phylogenetic methods proposed that retained marine dispersal abilities might explain transcontinental distributions in these lineages exclusively freshwater fishes (*51*), but are largely based on anecdotal evidence (*43*). Alternative hypotheses, such as the existence of island chains across the early Atlantic Ocean (*38*, *48*, *52*, *53*), have been proposed to explain how freshwater fishes and other terrestrial animals, such as dinosaurs, dispersed across oceans. The apparent conflict between the exclusively post-Cretaceous, freshwater fossil record and molecular divergence times of cichlids and other species-rich lineages such as killifishes (*24*, *27*, *38*, *48*, *54–57*) and their present-day distributions across continents that separated during the Mesozoic (*32*, *39*, *43*, *58*, *59*) has even been framed as a challenge to molecular clock methods used to calibrate the Tree of Life (*27*, *37*, *48*).

Here, we show that a single evolutionary mechanism accounts for both the exceptional species richness and present-day biogeography of cichlids and several other transcontinental freshwater fish radiations. Using a phylogeny of 591 cichlid species and representatives of candidate sister lineages of *Cichlidae*, we demonstrate that cichlids and several other exclusively freshwater fish lineages with transcontinental distributions originated in a rapid radiation near the Cretaceous-Paleogene boundary 66 million years ago. We show that saltwater tolerance, inherited from a common ancestor shared with species-rich marine fish lineages, explains the present-day distribution of cichlids and those of other closely related transcontinental freshwater clades, including killifishes, leaffishes, pupfishes, and livebearers. This retained ability to tolerate high salinities also explains how these lineages repeatedly colonized ephemeral inland waters, such as the African Rift Lakes and waterways of the western North American deserts, that have undergone salination and desiccation during the last few million years of global cooling. These results, which show how the retention of an ancestral physiological trait can enable lineages to access new ecological opportunities and subsequently diversify, reframes the importance of evolutionary context for understand how adaptive radiations assemble.

## Results

### A Cretaceous-Paleogene radiation unites transcontinental freshwater fish clades

The evolutionary relationships of cichlids among ray-finned fishes are historically controversial. Phylogenetic analyses consistently place cichlids in the clade *Ovalentaria*, which was first discovered in molecular phylogenetic analyses. Nevertheless, the identity of the successive sister lineages to *Cichlidae* remain unclear, in part owing to discrepancies in taxon sampling across studies (*27*, *60–65*).

We deployed a dataset of 1,314 ultraconserved element markers sequenced for 556 species in *Ovalentaria* and 35 outgroups using previously published sequences (*40*), available genomes, and newly-sequenced specimens from ovalentarian clades such as dottybacks (*Pseudochromidae*), eel blennies (*Congrogadidae*), and longfins (*Plesiopidae*). Despite this wealth of data, phylogenies generated using maximum likelihood and multispecies coalescent criteria across different taxon-sequence matrix completeness levels do not resolve consistent relationships among ten major lineages of *Ovalentaria*: *Atheriniformes* (killifishes, ricefishes, flying fishes, and silversides), *Embiotocidae* (surfperches), a clade containing *Blennioidei*, *Gramma*, *Opistognathidae*, and *Gobiesocidae* (blennies, clingfishes, and relatives), *Congrogadidae, Pomacentridae* (clownfishes and damselfishes), *Plesiopidae*, *Pseudochromidae*, *Polycentridae* (leaffishes), a clade composed of *Mugilidae* (mullets) and *Ambassidae* (Asiatic glassfishes), and a clade composed of cichlids and the eel-like engineer gobies *Pholidichthys* (Figure 1A, Figures S1-S7). Many of these lineages harbor species counts that are within one order of magnitude of cichlids (Figure 1B), suggesting that cichlid species richness alone is not wholly exceptional among closely related lineages of marine and freshwater fishes. This observation also implies that the evolutionary mechanisms facilitating cichlid species richness may be more widespread among lineages in *Ovalentaria*.

**Figure 1.**
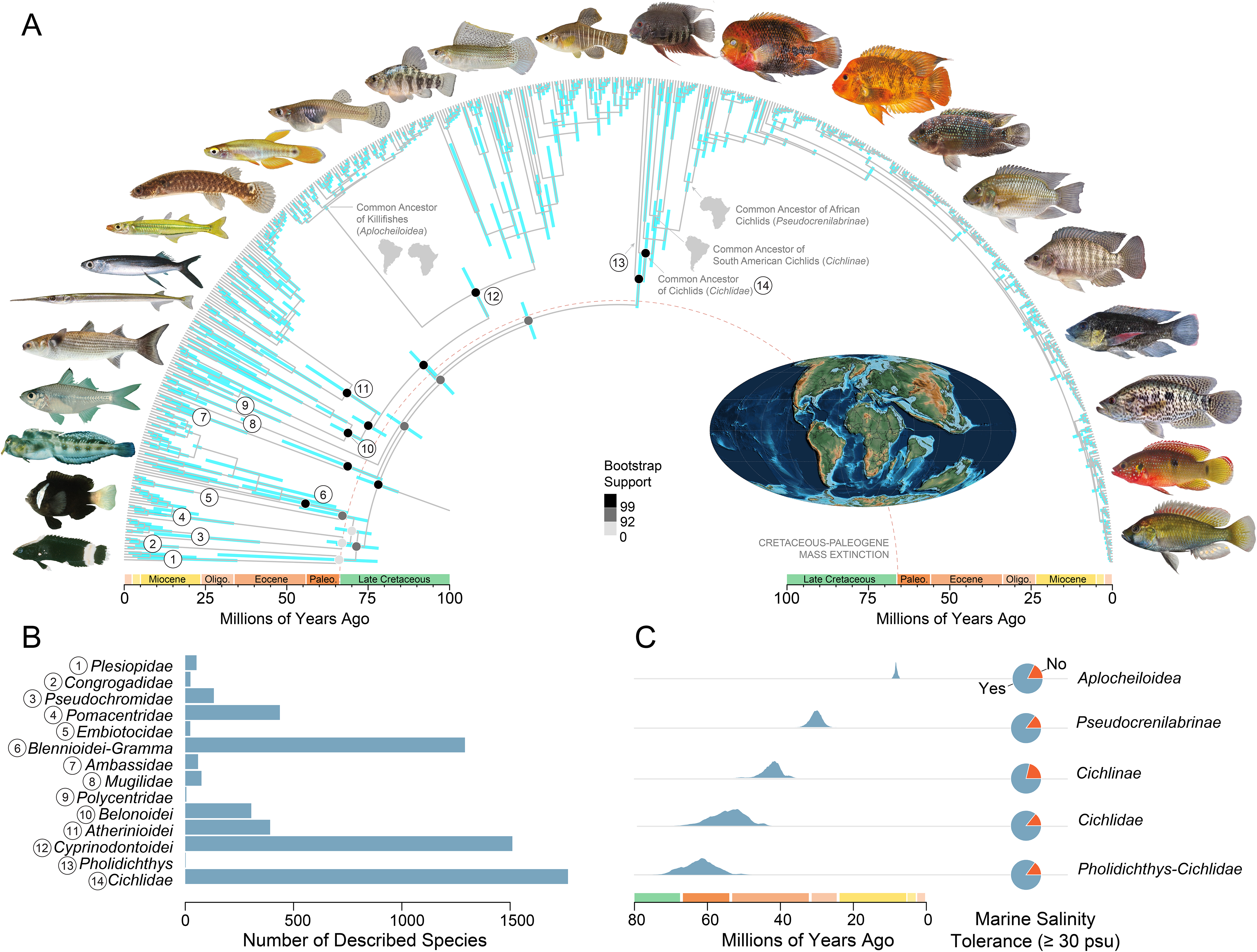
Phylogeny of *Ovalentaria* and the diversity of transcontinental freshwater radiations in context. (A) Time-calibrated phylogeny of *Ovalentaria* from Bayesian node-dating of three sets of 30 UCEs. Bars at nodes indicate 95% highest posterior density intervals of divergence times. Circles at nodes denote ultrafast bootstrap supports found in the single partition maximum likelihood phylogenetic analysis of the 90% complete dataset. Map is of the Earth 65 million years ago by Christopher Scotese (*172*), showing that all the southern continents had already separated by the time of the initial radiation of *Ovalentaria.* (B) Species counts across major lineages in *Ovalentaria* with similar crown ages from Eschemeyer’s Catalog of Fishes, accessed October 2025, showing that cichlids are within one order of magnidue of diversity of other clades of *Ovalentaria.* (C) Posterior distribution of MRCA ages across 20,000 randomly sampled trees from the posterior tree set (left) and inferred ancestral salinity tolerance presence (right) for selected transcontinental freshwater radiations. Note that *Cichlidae* and the ancestors of successive divergences between lineages restricted to particular continents (e.g., *Cichlinae* in the Americas, *Pseudocrenilabrinae* in Africa) are all inferred to have been tolerant of salinities ≥ 30 PSU. Photographs of fishes are by Zachary Randall (Florida Museum of Natural History, USA) and the late John E. Randall.

These uncertain relationships are suggestive of a rapid accumulation of diversity in *Ovalentaria* soon after its appearance. The successive divergences among the ten major lineages of *Ovalentaria* form a region of tree space called an anomaly zone, where more strongly supported alternative hypotheses of relationships among lineages to those in a species tree are found for a minority of gene trees (Figure S1) (*66*, *67*). Anomaly zones usually occur due to rapid successive lineage divergences, in which coalescence is rarely achieved across all genes (*66*, *68–71*). Despite the lack of resolution surrounding the relationships of clades in *Ovalentaria* (Figure 1, Figure S1), *Pholidichthys* is always inferred to be the sister lineage to *Cichlidae* with strong bootstrap support and with a comparatively high proportion of concordant sites (Figure S1). This result matches the resolution of cichlid relationships in previous taxon-rich phylogenies inferred using genome-wide marker data (*40*, *64*, *65*) and confirms that the closest living relatives of cichlids are marine fishes.

High discordance in lineage relationships often results from rapid successive divergences, such as those expected to occur during rapid evolutionary radiations (*67*, *72–76*). The uncertainty surrounding the phylogenetic relationships is consistent with a history of rapid origination for the major lineages of *Ovalentaria* (Figure S1) (*27*, *40*, *61–63*, *65*, *77*, *78*). Our time-calibrated phylogeny, based on 22 vetted fossils, demonstrates that the initial divergences among lineages of *Ovalentaria* all occurred around the Cretaceous-Paleogene boundary (Figure 1; Figure S1), establishing that cichlids are members of a larger rapid radiation triggered by the ecological restructuring following the Cretaceous-Paleogene Mass Extinction (*78*).

The posterior distribution of ages for cichlid lineages and the common ancestors of other exclusively freshwater transcontinental radiations in *Ovalentaria*, such as New World, African, and Asian killifishes (*Aplocheiloidea*), unambiguously reject vicariance induced by Mesozoic continental fragmentation as a mechanism driving their current distributions (Figure 1C; Extended Data Figures 2-5). In the new time-calibrated phylogeny that we generated, the age of *Ovalentaria* is estimated at 73.4 Ma, with 95% highest posterior density (HPD) intervals of 66.3116 and 80.034 Ma, the most recent ancestor (MRCA) of *Cichlidae* and *Pholidichthys* is estimated at 61.6093 Ma (95% HPD: 53.7374, 70.7134 Ma), the MRCA of *Cichlidae* is estimated at 53.6731 Ma (95% HPD: 44.2621, 62.6413 Ma), the MRCA of *Aplocheiloidea* is estimated at 8.3488 Ma (95% HPD: 7.4869, 9.1287 Ma), the MRCA of pupfishes and livebearers (*Cyprinodontoidea*) is estimated at 55.538 Ma (95% HPD: 46.9639, 63.9537 Ma), and the MRCA of rainbowfishes (*Melotaeniidae*) is estimated at 16.565 Ma (95% HPD: 10.9842, 21.2377 Ma). The age of *Aplocheiloidea* that we estimate is younger than the ages found in previous molecular phylogenetic analyses (*40*, *79*, *80*), but it is entirely consistent with the fossil record of this clade (*55*) and may be the result of our use of direct clade-specific age priors rather than outgroup calibration schemes. Our estimates for the age of *Cichlidae* and its constituent taxonomic subfamilies match or are slightly younger than those reported in prior phylogenomic analyses (*27*, *37*, *40*, *48*, *63*, *77*). The slightly younger ages that we estimate for the deepest divergences in cichlids are most probably attributable to the stem-ward placement of early African cichlid fossils following recent systematic revisions of extinct species (*24*). Our age estimates support the conclusion from previous time-calibrated phylogenies in which fossils, rather than continental fragmentation events, were used to place minimums on node ages (*26*, *27*, *37*, *38*, *40*, *48*, *62*, *63*, *77*, *78*, *81*) that cichlids, killifishes, and other transcontinental freshwater radiations in *Ovalentaria* originated and diversified long after the fragmentation of Mesozoic supercontinents.

### Ancestral physiology enabled transcontinental freshwater fish dispersals

Salinity tolerance provides an obvious candidate mechanism for explaining transoceanic dispersal in cichlids and other transcontinental freshwater fish radiations (*38*, *48*, *50*, *54*). Studies of cichlids and other lineages that exclusively inhabit fresh waters today have highlighted that lost legacies of saltwater tolerance might explain the present-day distribution of species (*44*, *46*, *48*, *50*, *82–85*). Our phylogeny confirms multiple lineages of transcontinental freshwater fishes in *Ovalentaria*, including *Aplocheiloidea*, *Cichlidae*, *Cyprinodontoidea*, and *Polycentridae,* all originated far after the separation of the southern continents (Figure 1, Extended Data Figures 2-5). The phylogenetic proximity of these lineages suggests a common mechanism such as salinity tolerance might explain the apparent mismatch between their geographic distribution and ages.

To test the hypothesis that retained tolerance to levels of salinity present in sea water explains the present-day biogeography of exclusively freshwater transcontinental radiations of fishes, we gathered data on tolerance (100% survival) to ≥ 30 practical salinity units (PSU), which is the lower end of salinity for ocean water (*86*, *87*), for species in our phylogeny. Our ancestral state reconstructions of salinity tolerance using data for 195 species under multiple models of character evolution supports the hypothesis that cichlids, aplocheiloid killifishes, and other transcontinental freshwater radiations all ancestrally retained tolerance to salinities characteristic of the marine realm (Figure 1, Figure S9). Salinity tolerance has been secondarily lost in clades endemic to particular continents, such as South and Central American cichlids, rainbowfishes, and *Nothobranchius* killifishes (Figure 1; Figure S9). These clades lost tolerance to marine salinities over an estimated 5 to 20 million years (Figure S9).Together, these results suggest that ancestral tolerance to marine salinities facilitated transoceanic dispersal in secondarily freshwater fish radiations with living species that have subsequently lost this physiological characteristic.

The retention of salinity tolerance in cichlids, killifishes, and other transcontinental radiations of fishes in *Ovalentaria* that we observe in our ancestral state reconstructions shows that physiological constraints which would otherwise restrict these fishes to fresh water did not exist when lineages endemic to different landmasses diverged (Figure 1). However, other phenotypic traits, such as body size, can also modulate marine dispersal ability(*88–90*). To examine the association of body size and habitat in *Ovalentaria*, we compiled measurements for n=169 species in our dataset (*91*) and conducted phylogenetic linear regressions of habitat occupancy against log-transformed proxies of body size. Neither body weight, length, depth, nor width are correlated with tolerance to ≥ 30 PSU or the occupancy of freshwater, brackish, and marine habitats by fishes in this lineage (Figure 2). Thus, body size is unlikely to have restricted the dispersal of freshwater fishes across oceans, nor is there evidence that smaller fishes are favored to undergo dispersal episodes in deep time (Figure 2).

**Figure 2.**
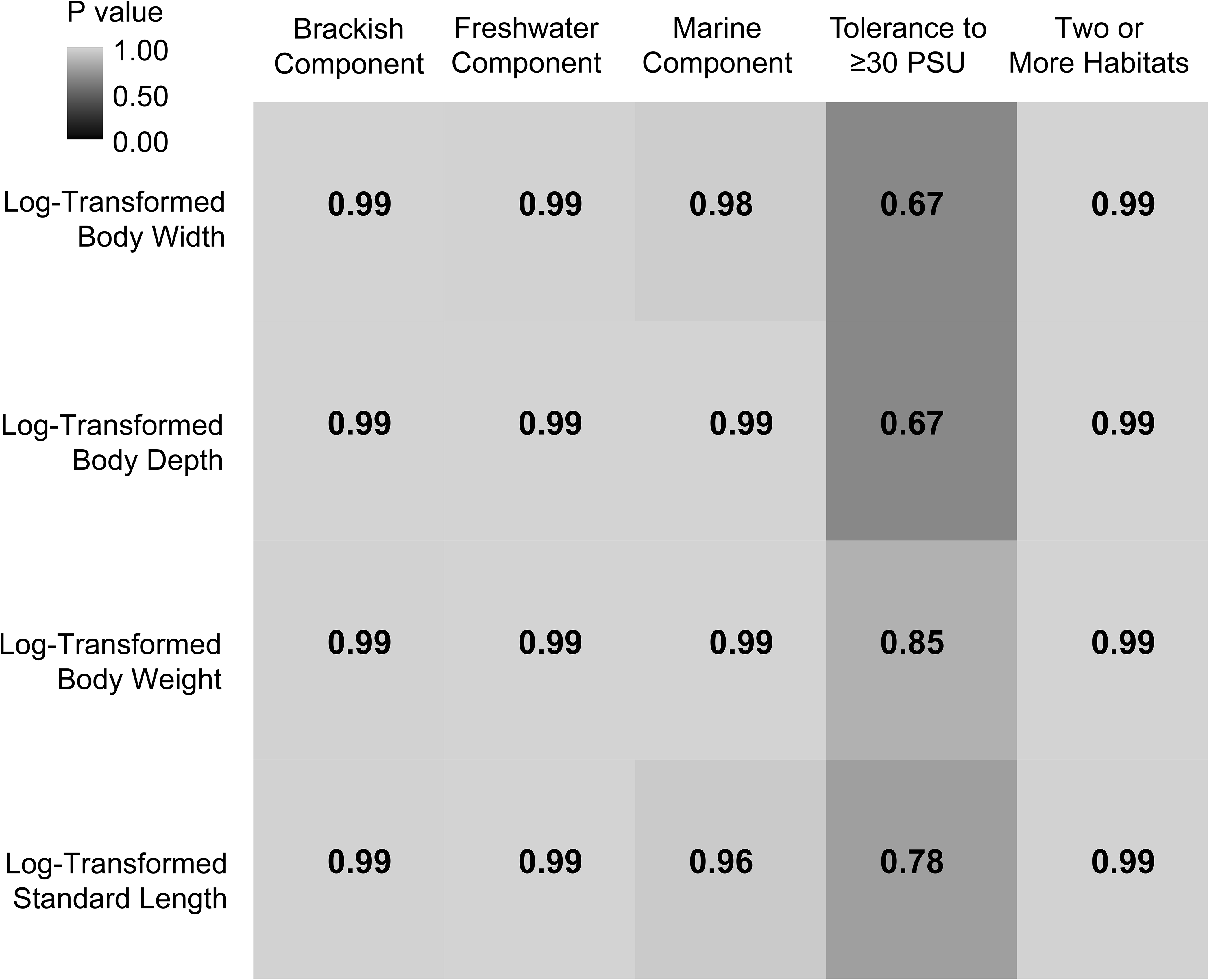
Body size is not associated with habitat in *Ovalentaria*. Heat map shows p values from phylogenetic logistic regressions of four body size proxies against binary habitat traits (freshwater/marine component to habitat) in the sample of n=168 species in *Ovalentaria*.

Ancestral state reconstructions of aquatic habitat occupancy support the persistence of brackish and saltwater habitat preferences deep into the major lineages in *Ovalentaria*, including *Cichlidae* and *Atheriniformes* (Figure 3A; Extended Data Figures 2-5). Our ancestral state reconstruction of habitat preference under a polymorphic character model estimates that the initial radiation of *Ovalentaria* took place in marine ecosystems, followed by multiple invasions into freshwater by species in *Ambassidae, Polycentridae, Adrianichthyidae*, *Aplocheiloidea*, *Cyprinodontoidea*, and the taxonomic subfamilies of cichlids (Figure 3A; Extended Data Figures 2-3). We reconstruct that a brackish-freshwater ecology is ancestral for the crown clades of the first two cichlid lineages to diverge: the Indo-Malagasy *Etroplinae* and *Ptychochrominae* (Figure 3A; Figure S4). A marine component is also reconstructed with ∼33% probability for the common ancestor of cichlids and the common ancestor of aplocheiloids and cyprinodontoids (Figure 3A). These results suggest that the initial diversification of these transcontinental radiations of fishes did not exclusively occur in freshwater environments. Our historical biogeographic reconstructions support the inference that saltwater tolerance and non-freshwater-exclusivity persisted deep into the major ovalentarian crown clades (Figure 3A). Across models, a dispersal-extinction-cladogenesis model with a jump dispersal parameter (DEC+j) was favored (Supplementary Information). However, all models supported a similar biogeographic scenario of a marine origin for *Ovalentaria* and highly uncertain areas of origin for the initial radiation of its ten major constituent lineages and the stepwise divergences of transcontinental lineages of cichlids, killifishes, and cyprinodontoids (Figure 3A). As expected based on their current distributions, the ancestral area of *Cichlinae* and *Rivulidae* is reconstructed as South America, the ancestral area of *Pseudocrenilabrinae* and *Nothobranchidae* is reconstructed as Africa, the ancestral area of *Aplocheilidae* and *Etroplinae* is reconstructed as Eurasia, and the ancestral area of *Ptychochrominae* is reconstructed as Madagascar (Figure 3A). We explicitly tested whether including saltwater tolerance has an effect on model fit using recently developed trait-mediated historical biogeographic reconstruction methods (*92*, *93*), but models incorporating trait-mediated biogeography did not improve fit relative to trait-free models (Supplementary Information). Because salinity tolerance is the ancestral condition in *Ovalentaria* and closely linked to the marine area state incorporated into our area delimitation (Figure 1, Figure 3), this trait may act as a redundant parameter in trait-mediated dispersal models.

**Figure 3.**
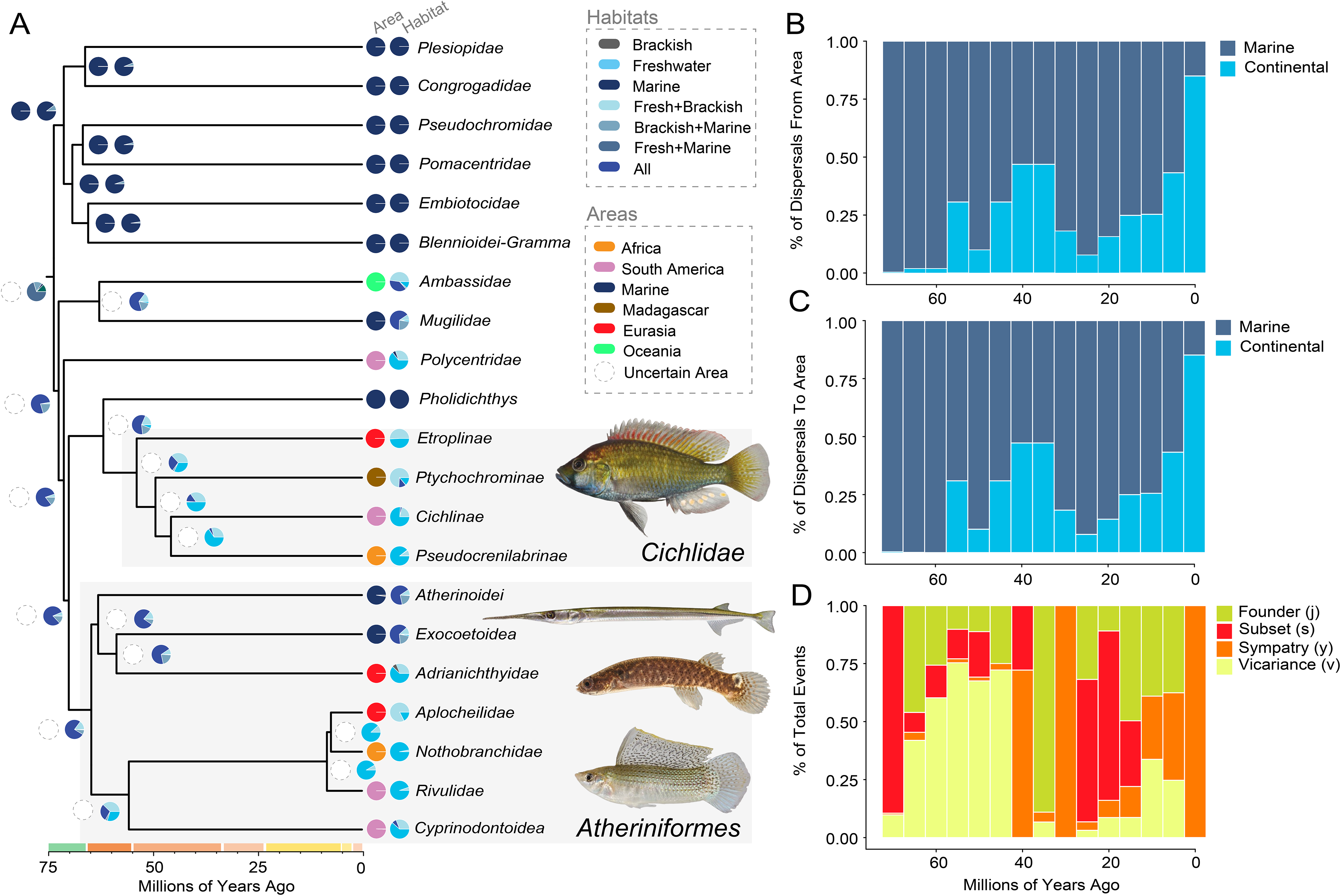
Historical biogeography and habitat suggests a lost legacy of marine inhabitation by transcontinental freshwater fish lineages. (A) Simplified phylogeny in Figure 1 showing the historical biogeographic reconstruction estimated under the best-fit model (DEC+j) in the full analysis, and the habitat ancestral state reconstruction estimated under a polymorphic character model in *phytools*. Highly uncertain nodes for the biogeographic reconstruction are denoted by open circles with dotted outlines. Photographs of fishes are by Zachary Randall (Florida Museum of Natural History, USA) and the late John E. Randall.

Marine-mediated dispersals across continents explain the uncertain location of the initial K-Pg radiation of *Ovalentaria* reconstructed in our analyses of historical biogeography. By running biogeographic stochastic mapping, we quantified the proportion of expected biogeographic events occurring at nodes through time. We reconstruct that the marine realm comprises the highest proportion of dispersal sources and destinations across included areas (Figure 3B-C; Figure S7). Dispersals to and from the marine realm are only outpaced by dispersals within continental settings over the last five million years, as expected given the recent accumulation of large proportions of species richness in exclusively freshwater-dwelling clades such as cichlids and killifishes (Figure 1, Figure 3A; Figure S7). The nature of the marine realm as the primary area of dispersal is also independent of the type of biogeographic event that is most common over any time slice (Figure 3D). Collectively, these results demonstrate that the ancestors of living transcontinental freshwater radiations in *Ovalentaria* were probably brackish and marine fishes whose tolerance to high salinities allowed them to disperse across oceans.

### Challenging habitats and functional innovations as diversification drivers

Although our ancestral state and historical biogeographic reconstructions demonstrate how cichlids and other lineages of *Ovalentaria* that are now restricted to fresh waters accessed new habitats across continents, these analyses do not directly examine how diversification regimes have been sculpted by biological traits like salinity tolerance. To evaluate how biological traits have affected lineage diversification in *Ovalentaria*, we estimated diversification rates associated habitat preference and salinity tolerance. We compared the effects of these traits on diversification to that of the fused pharyngeal jaw, the classic functional innovation in *Cichlidae*, *Pomacentridae*, and *Embiotocidae* which is hypothesized to enable ecological diversification by expanding feeding functionality (*60*, *65*, *94–96*). The number of distinct evolutionary origins of this feature within *Ovalentaria*, as well as its status as a diversification stimulant, are unclear (*65*, *97*, *98*). Our analyses of trait-associated diversification, including calculations of the DR statistic (*99*, *100*) and comparisons of Hidden State Speciation and Extinction (HiSSE) models (*101*), suggest that neither habitat preference nor the presence of a fused pharyngeal jaw is unambiguously associated with increased lineage diversification (Figure 4). Although the majority of fishes with a freshwater habitat component possess larger DR values than fishes that are exclusively brackish-marine (Figure 4A), diversification rate distributions estimated in our BiSSE analysis show that diversification rates associated with freshwater and marine habitat components do not significantly differ. The same pattern is present in our analysis in which the presence of a fused pharyngeal jaw is considered (Figure 4B). Character dependent hidden-state models best fit both trait sets, suggesting that neither habitat nor the presence of a fused pharyngeal jaw clearly explain the diversification history of *Ovalentaria* (Figure 4C).

**Figure 4.**
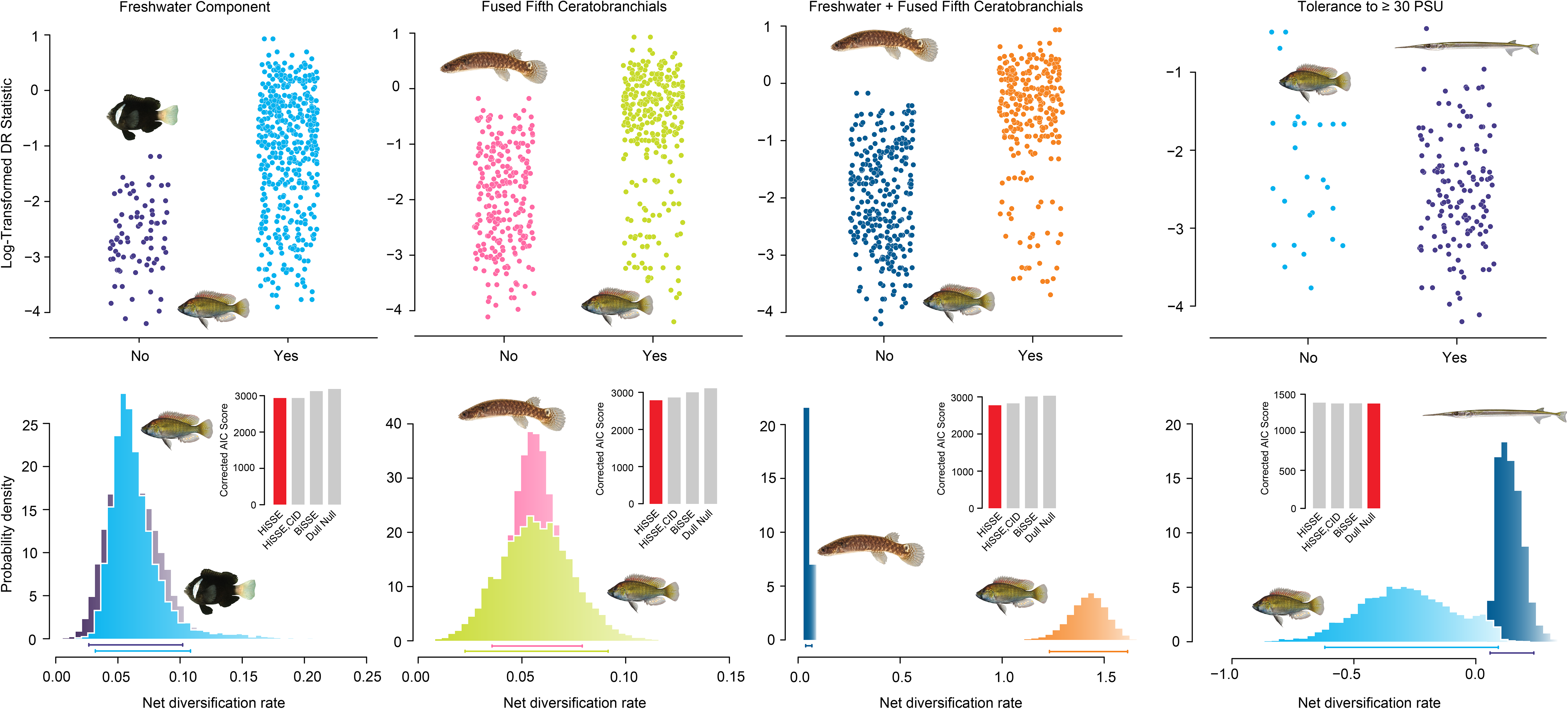
Indirect diversification effect of salinity tolerance. Plots of DR statistics by state (top) and BiSSE trait-associated diversification rate distributions (bottom) with inset model comparisons from the HiSSE analysis (bottom inset) for four binary traits: freshwater habitat component, marine habitat component, presence of fused pharyngeal jaws and a freshwater habitat component, and salinity tolerance. Bars underneath distributions in the lower half of the figure denote the 95% highest density interval for each trait-associated rate distribution.

When habitat preference and pharyngeal morphology are jointly considered, BiSSE analysis estimates that freshwater fishes with fused pharyngeal jaws exhibit higher diversification rates than other lineages in *Ovalentaria*, and a HiSSE character-dependent model is favored to explain differences in trait associated diversification rates among these groups. DR statistic values do not differ significantly between fishes that live in freshwater and possess fused pharyngeal jaws and those which do not possess this trait combination. Besides the common ancestor of *Cichlidae* and *Pholidichthys*, *Embiotocidae*, *Pomacentridae*, and a clade comprising flying fishes (*Exocoetidae*) and the paraphyletic *Hemirhamphidae* (halfbeaks) have independently evolved the fused pharyngeal jaw among *Ovalentaria* (Extended Data Figures 2-5) (*60*, *65*, *97*, *102*). Because all other lineages with a fused pharyngeal jaw, including *Pholidichthys*, are predominately marine, the combination of a freshwater habitat and fused pharyngeal jaw is unique to *Cichlidae*. Our results confirm that the elevated lineage diversification rate of cichlids is largely clade-specific rather than trait-driven (Figure 4, Figure S8), which is consistent with previous macroevolutionary analyses (*17*, *26*, *103*). Finally, we find no association between tolerance to marine salinities and elevated diversification across DR, BiSSE, and HiSSE analyses (Figure 4D). Thus, salinity tolerance cannot be considered a direct diversification catalyst in cichlids and other freshwater radiations of fishes in *Ovalentaria* (Figure 4D).

## Discussion

Sudden bursts of species richness are primarily attributed to two factors: ecological opportunity and innovation (*1–3*, *7*, *104*). This argument suggests that rapid lineage diversification is contingent on chance events, such as the evolution of new traits, the creation of new habitats, or the restructuring of ecosystems after a mass extinction (*1*, *2*, *105*). When the evolution of novel biological features coincide with the emergence of new habitats, their combined influence on diversification can be particularly powerful (*13*, *15*, *71*, *106*). Because of this perspective on how biodiversity accumulates, many analyses of evolutionary radiations focus on the lineages that rapidly diversify in isolation, without consideration of their macroevolutionary context (*8*, *11*, *17*, *26*, *107–110*). Yet, subsequent work on many adaptive radiations has shown that the key innovations (*65*, *71*, *102*) or ecological opportunity (*106*, *111*) proposed to trigger them are not supported as drivers of diversification when these systems are placed in a broader phylogenetic context. Many studies have also identified so-called pre-adaptations, which are features associated with rapid species diversification that predate its onset, across many celebrated examples of adaptive radiation (*9*, *17*, *25*). Together, these observations underscore the importance of broad macroevolutionary context for understanding what triggers evolutionary radiations.

Freshwater fish radiations, which are some of the most species-rich continental assemblages on the planet (*17*, *25*, *112*, *113*), include numerous lineages whose transcontinental biogeography appears at odds with their present-day physiology. The ongoing, century-long debate regarding the nature and prevalence of dispersal events in deep time largely originates from this observation (*43*, *46*, *83*, *85*, *88*, *114–116*). Our analyses provide a common explanation for these biogeographic patterns in several freshwater fish radiations whose timescale of evolution has been especially contentious, including cichlids and killifishes: salinity tolerance retained from their ancestry within radiation of marine fishes around the time of the Cretaceous-Paleogene boundary (Figure 1; Figure 3). This explanation is corroborated by phylogenetic analyses, ancestral state reconstructions, and historical biogeography, and is fully congruent with the fossil record of these lineages. Many lineages of *Ovalentaria* that our analyses show underwent transcontinental dispersals in the last 60 million years possess characteristics such as the retention of eggs inside the body (*80*, *117*) that could have increased the likelihood of successful colonization of a new habitat after a rare dispersal episode.

The retention of ancestral salinity tolerance across diverse freshwater lineages in *Ovalentaria* also provides a straightforward explanation for the concentration of rapidly diversifying lineages across the clade (Figure 1B). As originally conceived, key innovations are features that allow species to access new ecologies (*2*, *105*), and these features are not necessarily associated with a shift in diversification regime (*2*, *118*). Across the natural history of living freshwater species in *Ovalentaria*, there is widespread evidence that salinity tolerance has acted as a key innovation. *Cyprinodon* pupfishes show a variety of adaptations for salinity which allow them to tolerate isolated, hypersaline pools and rivers in the deserts of western North America (*119–121*), and hypersaline environments might drive phenotypic diversity in poecilid livebearers (*122*). Several cichlid species are now invasive to the Everglades along the southeastern coast of North America, where their dispersal ability has been directly modulated by this physiological feature (*123*, *124*). Even the cichlids that initially invaded Lake Victoria and gave rise to the exceptionally rapid radiation endemic to this water body (*6*) must have contended with a hypersaline and occasionally desiccated lake environment (*6*, *125*). Across these relatively young lineages, salinity tolerance has facilitated invasion and diversification in environments that might otherwise prove inhospitable. The ancestral state reconstructions that we present show that these recent radiations reflect a deeper history of physiology-mediated dispersal across *Ovalentaria* (Figures 1, 3).

The absence of a clear association between marine habitats, salinity tolerance, and diversification across *Ovalentaria* suggests that the effect of this physiological trait on the accumulation of species richness is more subtle than a direct causal relationship. Although our sample of species for which salinity tolerance has been documented is low relative to the total diversity of *Ovalentaria*, the data captures all major lineages in this clade. Rather than sampling bias, we suspect that our inability to detect a direct association between salinity tolerance and diversification is due to the exaptive (*126*) nature of this trait. The common ancestor of *Ovalentaria*, and likely all ray-finned fishes (*127–129*), was a marine animal, and so the effect of saltwater tolerance on diversification is theoretically already included in the background diversification rate of ray-finned fishes. A positive diversification effect of salinity tolerance would therefore only be detectable if the loss of this feature was associated with reduced diversification, but if salinity tolerance enabled fishes in *Ovalentaria* to disperse to new landmasses to radiate within freshwater ecosystems, we expect that salinity tolerance has enabled the rapid accumulation of species. This illustrates how a trait can simultaneously enable the invasion of new ecosystems and promote diversification without having a detectable effect on diversification rate.

How has the retention of salinity tolerance acted to trigger the rapid accumulation of diversity towards the present? The adaptive radiations of freshwater fishes such as cichlids, aplocheiloids, and cyprinodontoids have been attributed to a diverse set of traits and events in prehistory, including the presence of modified pharyngeal bones (*65*, *94*, *96*), lability in tooth complexity (*130*), the evolution of specialized feeding ecologies (*131*), and the origination of new habitats (*6*, *42*). The rapid diversification rate in cichlids, for example, shows high spatial dependence; it is almost entirely driven by the signal from African rift lake flocks (Figure S7) (*17*, *132*). The conclusion that increases in lineage diversification are driven by the unique parameters of novel habitats following invasions enabled by salinity tolerance is also supported by the fossil record of adaptive radiations. In the Triassic rift lakes of eastern North America, fishes in the extinct lineage †*Semionotidae* experienced replicated radiations that generated high morphological diversity, paralleling cichlid radiations today (*133*, *134*). The closest living relatives of semionotids are the species-depauperate and slowly-evolving gars (*135*, *136*), implying that the nature of rift lake ecosystems rather than the intrinsic biology of semionotids largely shaped the diversification of this lineage. Across these replicate radiations in similar geological contexts across deep time, the ability to disperse into new ecosystems provided the common ancestors of cichlids and other freshwater lineages with access to sources of ecological opportunity that differentially induced radiation.

The classic model of evolutionary radiation supposes the existence of ancestral biological features or an accessible source of ecological opportunity that subsequently spurs species diversification (*1*, *3*, *7*, *26*, *104*). An emerging consensus disputes this viewpoint and emphasizes the evolutionary (*71*, *111*, *130*, *137*) and ecological (*6*, *17*, *132*, *138*) context-dependence to the outcomes of ecological transitions and phenotypic novelties that precede rapid radiation. Here, we show that some of the most species-rich evolutionary radiations of freshwater fishes may have been enabled by a single retained ancestral feature: an ability to tolerate high salinities comparable to marine and coastal environments. This feature, along with a potentially recent history of marine inhabitation by living lineages exclusively found in freshwater, has propelled transcontinental dispersal in radiations such as cichlid fishes and aplocheiloid killifishes whose geographic distributions appear to reflect ancient vicariance induced by continental fragmentation. Because the ability to tolerate saltwater is ancestral for *Ovalentaria* and probably all ray-finned fishes (*129*), this feature is not clearly linked with increased diversification in freshwater radiations, but natural history observations support its importance as a mechanism for accessing new habitats and ecological opportunity. The revised evolutionary framework that we propose for cichlids, killifishes, and their relatives illuminates how ancestral physiological traits can shape the biogeographic and diversification legacies of entire clades across the Tree of Life.

## Methods

### Phylogenetic Rank-Free Taxonomy

In this manuscript, we deploy phylogenetic rank-free taxonomy following the protocols outline in the PhyloCode (*139*), which is now being applied to large swaths of living and extinct biodiversity (*140–142*), including ray-finned fishes (*143–145*). Following PhyloCode convention and emerging trends in the literature (*139*, *146*), we italicize all clade names in this manuscript. Our phylogenetic analyses of 1314 ultraconserved elements sequenced for 610 specimens representing 591 species of ray-finned fishes suggests that previous classification schemes for the clade *Ovalentaria* presented in phylogenetic rank-free taxonomies (*40*, *144*) are unreliable. *Blenniformes* as defined by Near and Thacker (*144*) is broadly paraphyletic across all of the phylogenomic analyses we present in this paper (Figure S1), as well as in several other recent studies of thousands of loci across the genomes of representative species in *Ovalentaria* (*27*, *63*, *65*, *77*). We propose a revision of this taxonomy where the lineages *Ambassidae*, *Blennioidei*, *Cichlidae*, *Congrogadidae*, *Embiotocidae*, *Gobiesocidae*, *Gramma*, *Lipogramma*, *Mugilidae*, *Opistognathidae*, *Pholidichthys*, *Plesiopidae*, *Polycentridae*, *Pomacentridae*, and *Pseudochromidae* are not placed in any order-level (i.e., -*iformes* group name) clade and are considered *Ovalentaria incertae sedis*. We highlight that this taxonomy is reflective of the highly uncertain relationships among *Ovalentaria* without necessitating the proliferation of new clade names for traditional order-and superfamily-level ranks that comprise single genera or families. For other rapid radiations, such as neoavian birds, this has led to redundancy between order, family, and genus names (e.g., *Struthioniformes*, *Struthionidae*, *Struthio*) which is, in our view, unnecessary at best and pedagogically harmful at worst. The clearest way forward that highlights the rapidity at which major lineages in *Ovalentaria* appeared around the K-Pg boundary is to keep the 15 clades noted above as unranked within any -*iformes* lineage.

### Sequence Dataset Assembly

We assembled sequences of the 1314 ultraconserved element loci (UCEs) from the Acanthomorphs 1k v.1 probe set (*78*) for a set of 610 specimens representing 591 species of ray-finned fishes to infer the phylogeny of *Ovalentaria* and test which lineage of ray-finned fishes is most closely related to *Cichlidae.* Previous studies that have investigated the phylogenetic relationships of *Cichlidae* have either incompletely sampled the family-level lineages of *Ovalentaria* (*27*, *63–65*, *147*), relied on small numbers of nuclear and mitochondrial markers (*60–62*, *148*), incompletely sampled the tribe-level lineages and radiations of *Cichlidae* and used single species to represent whole clades of *Ovalentaria* (*40*), or solely focused on cichlids (*26*, *32*, *149*). To simultaneously infer the interrelationships of *Ovalentaria* and all major lineages of cichlids, we combined previously published UCE data (*40*, *78*) with UCE sequences extracted from available genome assemblies and newly-target-sequenced specimens, including several new sequences for deeply-divergent lineages in *Congrogadidae*, *Plesiopidae*, and *Pseudochromidae*. We extracted, isolated, and prepared genomic DNA using previously published methods (*40*). All newly-isolated extractions were sequenced at the genomics facility of the University of Oregon, OR, USA.We used the program *phyluce* v. 1.7.3 (*150*) to process raw sequence reads, remove adapters, and remove low-quality bases via *Trimmomatic* (*151*) and *illumiprocessor* (https://github.com/faircloth-lab/illumiprocessor), assemble sequences, and remove duplicate UCEs. We also used *phyluce* v. 1.7.3 to harvest UCEs from published genomes and previously published read data, combine UCEs from these three sources (new sequences, previously published sequences, sequences skimmed from genomes), and align them to produce 75% and 90% complete taxon-character matrices. We reidentified one genome published on NCBI as “*Maratecoara gesmonei*” as a species of cichlid in the genus *Apistogramma*. We next used *CIAlign* (*152*) to visualize and check for chimaeric sequence alignments. Our final 75% and 90% complete UCE sets contained 1041 and 903 loci, respectively, for 594 specimens of *Ovalentaria* and 16 outgroups sampling all other taxonomic orders of *Acanthomorpha* (*144*). We conducted all bioinformatics processing steps on the Yale High Performance Computing Clusters (Yale HPC).

### Phylogenetic Analyses

We conducted maximum likelihood analyses on the 75% and 90% complete datasets using the phylogenetics program IQ-TREE v. 2 (*153*) on Yale HPC. For each dataset, we conducted three analyses: one in which we generated gene trees, one in which we concatenated UCEs and analyzed them under a single partition, and one in which we concatenated UCEs and analyzed them under a best-fit set of partitions found using PartitionFinder 2 (*154*). In each case, we used best-fit models of nucleotide sequence evolution found using ModelFinder (*155*) and calculated support for particular nodes using ultrafast bootstraps calculated over 1,000 replicates. For the gene tree sets, we inferred species trees under the multispecies coalescent model implemented in ASTRAL-III (*156*).

### Quantifying Phylogenetic Uncertainty and Anomaly Zones

To interrogate support for the phylogenies that we inferred beyond using ultrafast bootstrap supports, we calculated gene and site concordance factors, which measure the number of decisive gene trees and sequence sites that support a given relationship in an input species tree, using the ASTRAL-III multispecies coalescent topologies and the implementation of concordance factor calculation in IQ-TREE 2 (*157*). We also used custom scripts (*69*, *70*) to locate the position of anomaly zones along each species tree, which are nodes where a minority of gene trees contain stronger support for an alternative set of relationships to that present in the species tree (*67*). We only considered nodes where an alternative topology was favored by over 20 gene trees to fall in anomaly zones.

### Time-Calibration

In order to test whether Mesozoic continental fragmentation could conceivably have generated the present-day distributions of *Cichlidae*, *Aplocheiloidea*, and other transcontinental radiations of exclusively freshwater fishes, we assembled a dataset of 21 justified fossil calibrations (Supplementary Information) to calibrate ingroup nodes across *Ovalentaria* and the MRCAs of several outgroups. We used these fossil calibrations and three randomly subsampled sets of 30 UCEs to estimate a time-calibrated phylogeny in BEAST v. 2.6.7 (164, 165) using the BEAST implementation of the Fossilized Birth-Death Model (166). For each subset, we used a Lognormal Relaxed Clock Model and a General Time Reversal model of nucleotide evolution with the Gamma rate heterogeneity parameter. We set the value of rho, the number of living species sampled, to 0.1, which is the approximate number of species in our dataset divided by the number of species in Ovalentaria listed on Eschmeyer’s Catalog of Fishes in October of 2025 (167). We set the diversification rate prior to 0.05, which is the approximate background rate in acanthomorphs (41), the prior on the origin to 145.0 Ma. We used the age of each fossil calibration to sculpt log-normal MRCA priors such that 97.5% of the prior distribution fell before the age of the fossil. We fixed the topology to the maximum likelihood single-partition concatenated tree inferred using the 90% complete dataset. For each run, we ran Markov Chain Monte Carlo chains for 2.0 x 10^8^ generations with a 2.0 x 10^8^ generation pre-burnin. We pooled posterior log files for each independent run and checked for effective sample size values greater than 200 and convergence of posteriors in Tracer v. 1.7 (*158*), combined posterior tree sets using LogCombiner v. 2.7.7 (*159*), and annotated divergence times to the target tree in TreeAnnotator v. 2.6.7 (*159*)

### Phylogenetic Logistic Regressions

To assess for relationships between body size proxies and habitat, we conducted phylogenetic logistic regressions of the presence of a marine habitat component and four proxies of body size using the *phyloglm* function in the R package *phylolm* v. 2.1 (*160*): log-transformed standard length, log-transformed body weight, log-transformed body depth, and log-transformed body width. These measurements were taken for a subsample of n=168 species in our dataset present in the FishShapes v. 1 database (*91*).

### Discrete Trait Ancestral State Reconstructions and Evolutionary Lag-Times

To reconstruct the evolution of salinity tolerance, habitat state, and fused pharyngeal jaw presence across *Ovalentaria*, we conducted ancestral state reconstructions using the R package *phytools* v. 2.0 (*161*). We collected information on tolerance to ≥ 30 PSU salinity for n=194 species in our phylogeny and the presence of the fused pharyngeal jaw for all species in our phylogeny from the literature, as well as information on habitat (freshwater-brackish-marine) for all species in our phylogeny by searching FishBase (*162*). We differentially treated the habitat data as either a binary character coded by the presence of a freshwater or marine component or as a polymorphic character. For all binary character, we assessed the fit of four models of character evolution: one in which transition rates between state were fixed as equal (ER), one in which all transition rates differed (ARD), a model wherein reversals to the zero state were forbidden, and finally a model wherein reversals to the one state were forbidden. We compared the fit of these models using Akaike Information Criterion (AIC) scores and selected the model with the lowest score for further analysis. For the tolerance dataset, we decided to conduct ancestral state reconstructions using both the best-fit ARD model and the first of the two irreversible models, since salinity tolerance might be a complex trait that at least partially matches expectations of Dollo’s Law of Irreversible trait evolution. For the analysis where we treated habitat as a polymorphic character, we used the *fitpolyMk* function, an ARD transition rate model, and the root prior distribution π from FitzJohn et al. (*163*) to construct the transition matrix. For all analyses, we ran stochastic mapping over 1,000 simulated topologies and summarized them in single ancestral state reconstructions.

Our analyses suggest that salinity tolerance has been lost in a number of freshwater fish radiations in *Ovalentaria* endemic to particular continents. To understand the timescale of salinity tolerance loss following continental invasion, we wrote custom scripts with the aid of *Claude* Opus 4.7 to collect the ages of nodes where salinity tolerance was lost in our ancestral state reconstruction, and the nodes immediately subtending these, across 20,000 trees randomly selected from our posterior time tree set. Next, we used *ggplot2* (*164*) to produce histograms and density curves for these ages to represent the probability densities of key innovation origins and swim bladder losses through time.

### Historical Biogeography

*Trait-free.* We reconstructed the historical biogeography of *Ovalentaria* using our time-calibrated phylogeny and geographic data taken from FishBase using the R package *BioGeoBEARS* (*114*). For the input areas, we used the following: Americas (North, Central, and South America, and the Caribbean islands), Africa, Eurasia (to Wallace’s line), Oceania (to Wallace’s line), Madagascar, and finally a ‘marine realm’ area. Using AIC scores, we compared the fit of three alternative models of historical biogeography with and without the jump dispersal parameter j: a dispersal-extinction-cladogenesis (DEC) model, a dispersal-vicariance-like (DIVALIKE) model, and a Bayesian model (BAYAREALIKE). In addition, we reconstructed historical biogeography using all models to assess for major differences in estimated historical biogeographic reconstructions across model types. After the best-fit model was selected, we re-ran historical biogeographic reconstruction using biogeographic stochastic mapping (BSM) analysis. This analysis estimates the type of biogeographic events and their frequency through time by simulating histories (*165*). We used 100 as the maximum number of maps to try, 50 as the goal number, and 400 tries per branch. We extracted and plotted the number of cladogenetic events estimated from the output by region and by event type.

### Trait-mediated

Biological traits can affect the ability of species to disperse across areas, and therefore including a discrete trait parameter in historical biogeographic reconstruction might modify estimated biogeographic histories. Using recently developed code in *BioGeoBEARS* (*93*), we assessed the fit of trait-mediated and trait-independent biogeography models in which salinity tolerance was the discrete evolving trait considered to mediate dispersal. The inclusion of an evolving trait adds both trait state forward and reverse transition parameters (*t*_12_ and *t*_21_), as well as multipliers on dispersal probability based on a lineage being in trait state one (*m*_1_) or state two (*m*_2_) when a binary trait is considered. Using the pruned taxon set and corresponding geography data, we tested whether the model type (DEC/DEC+j) that best fit the data in the full analysis outperformed the same basic biogeographic model with the addition of these transition rate parameters and multipliers.

### Diversification Analyses

#### DR Statistic

To test for the influence of habitat, salinity tolerance, and the fusion of the pharyngeal jaw on diversification across *Ovalentaria*, we deployed several approaches for estimating trait-mediate diversification. First, we used the DR statistic, a measurement of diversification calculated for each tip in the phylogeny that accounts for distances between nodes and the number of successive branching events that occur between tips and the tree root (*166*), using the *EcoPhyloMapper (epm)* v. 1.1.6 package in R (*167*). Because of how it is calculated, the DR statistic is best understood as a metric of speciation rate towards the presence (*166*). Following previous studies (*106*, *168*), we calculated DR statistics for all species of *Ovalentaria* in our phylogeny after removing outgroups, and then compared the distributions of DR statistics for species grouped by habitat, the presence of the fused pharyngeal jaw, and salinity tolerance using phylogenetic generalized least squares (PGLS) implemented in the R package *nlme* (*169*). We also plotted DR statistics per major lineage of *Ovalentaria* to compare diversification rates of *Cichlidae* and other species-rich lineages in this clade (*17*).

#### Trait-Associated Diversification Rates

We deployed a binary state speciation and extinction model (BiSSE) to test for differences between the four binary traits (pharyngeal jaw fusion, presence of a freshwater component to habitat, presence of a marine component to habitat, and tolerance to ≥ 30 PSU, using the R package *diversitree* (*170*). After testing likelihood models where speciation rates were alternatively forced to be equal or allowed to differ across states, we ran MCMC chains for 10,000 generations on the best fit model in each case, sampling every 100 generations. After confirming convergence of the posteriors following burning in 10% of generations, we plotted histograms of estimated diversification rates associated with each state. Next, we used the R package *hisse* v. 2.1.11 (*171*) to assess the fit of trait-associated diversification rate models that alternatively included hidden states. We assessed the fit of four models: a ‘dull null’ model in which the turnover and extinction fractions were the same across all states, a BiSSE model in which turnover and extinction fractions were allowed to between lineages that possessed or lacked a swim bladder, a character-dependent hidden state speciation and extinction (HiSSE CD) model in which turnover and extinction fractions were allowed to vary between both lineages that possessed or lacked a swim bladder *and* between hidden states introduced into the analysis, and finally a character-independent hidden state speciation and extinction (HiSSE CID) model where turnover and extinction fractions were allowed to vary across hidden states, but not across known states. We compared model fits using AIC scores. For all diversification rate analyses, we removed outgroups and input sampling fractions by dividing the number of species known to possess a particular trait state in our dataset by the total known across *Ovalentaria*. Because we did not know the total number of salinity-tolerant species across *Ovalentaria*, we set the sampling fraction to 0.1 in that analysis.

In addition to trait-dependent speciation models, the R package *hisse* also allows for the use of a missing state speciation and extinction model (MiSSE) whereby only hidden states govern diversification regimes across the phylogeny (*101*, *171*). We used this to estimate diversification rates across our phylogeny and assess for differences in diversification across lineages. After using the *generateMiSSEGreedyCombinations* with a maximum of four parameters to construct MiSSE models, we used *MiSSEGreedy* to execute missing state models along our phylogeny. We input a value of 0.1, the approximate number of species in our dataset divided by the number of species in *Ovalentaria* listed on Eschmeyer’s Catalog of Fishes in October of 2025, as the sampling fraction. After comparing the fit of these models, we plotted the diversification rate phylogram found in the best-fit model.

## Acknowledgements.

We thank Nicholas J. Matzke and Sarah-Sophie Weil for help related to trait-mediated biogeographic reconstruction analysis. We thank Gregory J. Watkins-Colwell for help with specimen acquisition and handling at the Yale Peabody Museum, and the ASU Fish and Tissue Collection for loaning tissues. We finally wish to thank Zachary Randall (Florida Museum of Natural History, USA) and the Bernice Pauahi Bishop Museum and late John E. Randall for high-resolution photographs of representative species used in our figures.

## AI Statement

No AI technologies were used in the writing of this paper.

## Data Availability

All data from this study is either available in the Supplementary Information associated with this article or the associated Yale Dataverse Repository (https://doi.org/10.60600/YU/OQDK2P). New sequences will be uploaded to NCBI GenBank sequence archive under BioProject number XXX.

