## Supplemental Text, Tables, and Figures for "Physiology mediates transcontinental radiations of freshwater fishes"

**Table of Contents:**

Fossil Calibration Justification List.

Supplementary References.

Supplementary Figures and Captions.

Supplementary Tables

**Fossil Calibration Justification List.**

†*Cumbaaichthys oxyrhynchus*

**Phylogenetic Placement and Justification:** †*Cumbaaichthys oxyrhynchus* calibrates pan-*Polymixia*; in analyses presented in this paper, this is the MRCA of *Polymixia lowei* and *Chologaster cornuta*. See a full justification of this calibration in [1].

**Stratigraphy**: Lacdes Bois, Northwest Territories, Canada: early Turonian, Late Cretaceous [1].

**Fossil tip age**: 93.9 Ma [2,3].

†*Trichophanes foliarum*

**Phylogenetic Placement and Justification:** †*Trichophanes foliarum* calibrates pan-*Aphredoderus*; in analyses presented in this paper, this is the MRCA of *Chologaster cornuta* and *Percopsis omiscomaycus*. This placement is based on phylogenetic analyses of morphological characters [4]. See a full justification of this calibration in [5,6].

**Stratigraphy**: Florissant Fossil Beds, Colorado, USA: Priabonian, Eocene 37.7-33.9 Ma [6].

**Fossil tip age**: 35.8 Ma.

†*Iridopristis parrisi*

**Phylogenetic Placement and Justification:** †*Iridopristis parrisi* calibrates the MRCA of *Beryx splendens* and *Holocentrus rufus* and is placed as a pan-holocentrid in a recent phylogenetic analysis of morphological characters [7].

**Stratigraphy:** Main Fossiliferous Layer, Hornerstown Formation, New Jersey, USA: earliest Danian, Paleocene [7], approximately 66.02 Ma [8].

**Fossil age**: 66.02 Ma.

†*Gasterorhamphosus zuppichini*

**Phylogenetic Placement and Justification:** †*Gasterorhamphosus zuppichini* calibrates the MRCA of *Thunnus orientalis* and *Fistularia commersonii*. †*G. zuppichini* is the oldest identifiable member of *Syngnathiformes* and is placed in the crown of that order based on the results of phylogenetic analyses of morphological characters [9].

**Stratigraphy**: Nardo, Italy; Late Cretaceous, Santonian-Campanian, 83.6 Ma [9].

**Fossil tip age**: 83.6 Ma.

†*Ctenoplectus williamsi*

**Node calibrated:** †*Ctenoplectus williamsi* is a stem-triodontid in *Tetraodontoidei* that calibrates the node representing the MRCA of the clade containing *Triacanthodes ethiops* and *Siganus spinus* (*Acanthuriformes*) and is placed sister to *Triacanthodes ethiops* in the tip-dating analyses. The identity of †*Ctenoplectus williamsi* as a member of *Triodontidae* is supported by: jaws form acute arrowhead shapes in dorsal and ventral views, dental microstructure consists of simple, round elements densely stacked together, the presence of osteodentinic posterointernal walls and absence of anteroexternal walls in the dental microstructure, and the presence of posttemporals, ribs, pelves, and spines in the dorsal fin [10].

**Stratigraphy**: London Clay Formation, England, UK: Ypresian, Eocene, Paleogene 53 ma [10].

**Fossil tip age**: 53.0 Ma. However, the node calibration was created such that 97.5% of the probability distribution fell before 56.0 Ma given that multiple species of tetraodontiform of unstable affinity are known from approximately that age and this is the upper bound of the age of †*Ctenoplectus* [10].

†*Mene purydi*

**Phylogenetic Placement and Justification:** †*Mene purydi* is assignable to pan-*Mene* and calibrates the MRCA of *Betta prima* and *Caranx caninus* in our phylogeny. See a full justification of this calibration in [5,6].

**Stratigraphy**: Talara Province, northwestern Perú. Thanetian to Ypresian, Eocene [11], minimum 56.0 Ma [8].

**Fossil tip age**: 56.0 Ma.

†*Mioblennius fraudulentus*

**Phylogenetic Placement and Justification:** †*Mioblennius fraudulentus* is assignable to *Blenniidae* based on skeletal anatomy, but cannot be confidently placed within the crown [12]; as such, it calibrates the MRCA of *Salarias fasciatus* and *Gillellus semicinctus* in our phylogeny.

**Stratigraphy**: Spirialis Clays, Azerbaijan; Chokrakian regional stage of the Middle Miocene [12], 14.8 to 13.8 Ma [13].

**Fossil tip age**: 14.75 Ma.

†*Clinitrachoides gratus*

**Phylogenetic Placement and Justification:** †*Clinitrachoides gratus* is assignable to *Clinidae* based on skeletal anatomy, but cannot be confidently placed within the crown [12,14]; as such, it calibrates the MRCA of *Labrisomus nuchipinnis* and *Gibbonsia metzi* in our phylogeny.

**Stratigraphy**: Naslavcha, Karpov Yar, Moldova [14]; Sarmatian regional stage (upper Serravallian) of Miocene; 12.2-11.6 Ma [15,16].

**Fossil tip age**: 11.6 Ma.

†*Mugil princeps*

**Phylogenetic Placement and Justification:** †*Mugil princeps* is assignable to *Mugilidae* based on skeletal anatomy [17,18] and calibrates the MRCA of *Mugil cephalus* and *Chelon labrosus* in our phylogeny.

**Stratigraphy**: Menilite Shales, Ukraine; Rupelian Stage of the Oligocene [8], 33.9 to 27.3 Ma [8].

**Fossil tip age**: 27.3 Ma.

†*Hemitrichas stapfi*

**Phylogenetic Placement and Justification:** †*Hemitrichas stapfi* is assignable to *Atherinidae* based on its high vertebral count and otolith morphology [19] and calibrates the MRCA of *Atherinomorus stipes*, *Atherina presbyter*, and *Craterocephalus stercusmuscarum*.

**Stratigraphy**: Upper portion of the lower Ceritherium Beds, Dexheim, Germany; latest Oligocene, minimum age uppermost Oligocene, 23.0 Ma [20].

**Fossil tip age**: 23.0 Ma.

*Odontesthes* sp.

**Phylogenetic Placement and Justification:** *Odontesthes* sp. is assignable to Pan-*Odontesthes* based on the presence of a fenestra on the opercle and an unrestricted protractile premaxilla [19,21].

**Stratigraphy**: Ñirihuau Formation, Chubut Province, Argentina; early Miocene, 20.0 Ma [21].

**Fossil tip age**: 20.0 Ma.

†*Atherina atropatiensis*

**Phylogenetic Placement and Justification:** †*Atherina atropatiensis* is assignable to crown *Atherina* based on comparative meristic counts and skeletal anatomy [22] and calibrates the MRCA of *Atherina boyeri* and *A. presbyter* in our phylogeny. †*A. atropatiensis* most closely resembles *A. boyeri* in meristic counts and simplified vertebral anatomy [22].

**Stratigraphy**: Lignite Beds, Tabriz Basin, Iran; middle Miocene, 11.6-10.0 Ma [23].

**Fossil tip age**: 10.0 Ma.

†*Kenyaichthys kipkechi*

**Phylogenetic Placement and Justification:** †*Kenyaichthys kipkechi* is assignable to *Aplocheiloidei* as a member of pan-*Rivulidae* based on a phylogenetic analysis of morphological characters [24] and calibrates the MRCA of *Atlantirivulus santensis* and *Aplocheilus panchax* in our phylogeny.

**Stratigraphy**: Lukeino Formation, Tugen Hills, Kenya; late Micoene, 6.0-5.7 Ma [24,25]

**Fossil tip age**: 5.7 Ma.

†*Rhamphexocoetus evolans*

**Phylogenetic Placement and Justification:** †*Rhamphexocoetus evolans* is assignable to *Belonoidei* [26] sensu Near and Thacker [6] and calibrates the MRCA of *Oxyporhamphus micropterus* and *Xenentodon cancila* in our analysis. The conservative placement of this fossil outside of the crown of *Exocoetidae* + *Hemirhamphidae* is due to the presence of numerous features that exclude †*R. evolans* from both families [26].

**Stratigraphy**: Monte Bolca, near Verona, Italy: early-late Ypresian, Eocene, Paleogene 48.96 to 48.5 Ma [27].

**Fossil tip age**: 48.5 Ma.

†*Empetrichthys erdisi*

**Phylogenetic Placement and Justification:** *Empetrichthys erdisi* is assignable to *Goodeidae* as a member of Pan-*Empetrichthys* based on the absence of a pelvic girdle and fins, presence of a few rows of conical teeth, posteriorly placed dorsal and anal fins, and meristic counts [28], and calibrates the MRCA of *Crenichthys baileyi* and *Ameca splendens* in our analysis.

**Stratigraphy**: Posey Canyon shale, Peace Valley Formation, California, USA; Late Miocene [29,30], 8.4-5.0 Ma.

**Fossil tip age**: 5.0 Ma.

†*Fundulus lariversi*

**Phylogenetic Placement and Justification:** *Fundulus lariversi* is assignable to *Fundulus* and subgenus *Fundulus* based on a phylogenetic analysis of morphological and molecular data [31] and calibrates the MRCA of *F. heteroclitus* and *F. catenatus* in our analysis [30,31].

**Stratigraphy**: Siebert Tuff, Tonopah, Nevada, USA; Early to Middle Miocene, 17.8 to 16.2 Ma [31].

**Fossil tip age**: 16.2 Ma.

†*Prolebias stenoura*

**Phylogenetic Placement and Justification:** †*Prolebias stenoura* is assignable to Pan-*Valenciidae* based on a phylogenetic analysis of morphological characters [32] and calibrates the MRCA of *Apricaphanius iberus* and *Valencia letourneuxi* in our node-dating analyses.

**Stratigraphy**: Corent, Puy-de-Dôme, France; Rupelian Stage of the Oligocene [33], 33.9-28.1 Ma [8].

**Fossil tip age**: 28.1 Ma.

†*Chaychanus gonzalezorum*

**Phylogenetic Placement and Justification:** †*Chaychanus gonzalezorum* is assignable to Pan-*Pomacentridae* [34] and calibrates the MRCA of *Pomacentrus philippinus* and *Cypho purpurascens* in our node-dating analyses.

**Stratigraphy**: Belisario Domínguez quarry, Tenejapa-Lacandón geological unit, Chiapas, Mexico; Danian, 63.0 ± 1.5 Ma [34].

**Fossil tip age**: 63.0 Ma.

†*Plesioheros chauliodus*

**Phylogenetic Placement and Justification:** †*Plesioheros chauliodus* is assignable to Pan-*Heroini* [35] and calibrates the MRCA of *Pterophyllum scalare* and *Cichlasoma orientale* in our node-dating analysis.
**Stratigraphy**: Faja Verde, Lumbrera Formation, Alemanía, Argentina; Eocene, 39.9 Ma [35].
**Fossil tip age**: 39.9 Ma.

†*Libyachromis fugacior*
**Phylogenetic Placement and Justification:** †*Libyachromis fugacior* is assignable to Pan-*Pseudocrenilabrinae* based on phylogenetic analyses of morphological characters [36,37] and calibrates the MRCA of *Heterochromis multidens* and *Rhamphochromis esox* in our node-dating analysis. Murray [36] recently highlighted the problematic assignment of the older (45-46 Ma) †*Mahengechromis* spp., which is often used as a calibration for Pan-*Pseudocrenilabrinae* [25,38,39], to that clade. As such, we here use the earliest Oligocene occurrences of definite stem lineages of *Pseudocrenilabrinae*.
**Stratigraphy**: Taráb Formation at Jabal Al Hasáwnah, northeast of Trúnzah Hill near Wádi Taráb, Libya; Rupelian Stage of the Oligocene [37], 33.9 to 27.3 Ma [8].
**Fossil tip age**: 27.3 Ma.

†*Oreochromimos kabchorensis*
**Phylogenetic Placement and Justification:** †*Oreochromimos kabchorensis* is assignable to Pan-*Oreochromini* [40] and calibrates the MRCA of *Oreochromis niloticus* and *Coptodon congicus* in our node-dating analysis.
**Stratigraphy**: Kabchore, site 2a and 2b, Member C, Ngorora Formation, Tugen Hills, Kenya; Miocene, 12.5 Ma [40].
**Fossil tip age**: 12.5 Ma.

†*Tugenchromis pickfordi*
**Phylogenetic Placement and Justification:** †*Tugenchromis pickfordi* is assignable to the total clade of the East African Radiation based on phylogenetic analyses of morphological characters [36,37,41] and calibrates the MRCA of *Tilapia sparmanii* and *Bathybates ferox* in our node-dating analysis.

**Stratigraphy**: Outcrop Waril, Member E of the Ngorora Formation, Kenya; Late Miocene, 9-10 Ma [41].

**Fossil tip age**: 9 Ma.

**References.**

1. Murray, A. Mid-Cretaceous acanthomorph fishes with the description of a new species from the Turonian of Lac des Bois, Northwest Territories, Canada. *Vertebr. Anat. Morphol. Palaeontol.* **1**, 101–115 (2016).
2. Wilson, M. V. H. A Second Species of Libotonius (Pisces: Percopsidae) from the Eocene of Washington State. *Copeia* **1979**, 400–405 (1979).
3. Mayr, G., Archibald, S. B., Kaiser, G. W. & Mathewes, R. W. Early Eocene (Ypresian) birds from the Okanagan Highlands, British Columbia (Canada) and Washington State (USA). *Can. J. Earth Sci.* **56**, 803–813 (2019).
4. Murray, A. M., Brinkman, D. B., Newbrey, M. G. & Neuman, A. G. Earliest North American articulated freshwater acanthomorph fish (Teleostei: Percopsiformes) from Upper Cretaceous deposits of Alberta, Canada. *Geol. Mag.* **157**, 1087–1096 (2020).
5. Ghezelayagh, A. *et al.* Prolonged morphological expansion of spiny-rayed fishes following the end-Cretaceous. *Nat. Ecol. Evol.* **6**, 1211–1220 (2022).
6. Near, T. J. & Thacker, C. E. Phylogenetic classification of living and fossil ray-finned fishes (Actinopterygii). *Bull. Peabody Mus. Nat. Hist.* **65**, 3–302 (2024).
7. Andrews, J. V., Schein, J. P. & Friedman, M. An earliest Paleocene squirrelfish (Teleostei: Beryciformes: Holocentroidea) and its bearing on the timescale of holocentroid evolution. *J. Syst. Palaeontol.* **21**, 2168571 (2023).
8. Gradstein, F. M., Ogg, J. G., Schmitz, M. & Ogg, G. *The Geologic Time Scale 2020*. (Elsevier Science, Amsterdam, The Netherlands, 2021).

**Figure S1. Phylogenetic uncertainty during the initial rapid radiation of *Ovalentaria.*** (A) simplified phylogenies from each of the six different analyses across two levels of taxon-character matrix completeness, showing the highly uncertain relationships among ten major lineages of *Ovalentaria*. Grey bars highlight anomaly zone at the base of *Ovalentaria*. (B) Heat map showing coalescent support, ultrafast bootstrap support, and gene and site concordance factor values for selected nodes representing the initial divergences among lineages of *Ovalentaria*. Note that the monophyly of a clade containing *Cichlidae* and *Pholidichthys* is strongly supported across all analyses.

**Figure S2.** ASTRAL-III multispecies coalescent phylogeny inferred using the 75% complete matrix.

**Figure S3.** Maximum likelihood phylogeny inferred using the concatenated 75% complete matrix treated as one partition.

**Figure S4.** Maximum likelihood phylogeny inferred using the concatenated 75% complete matrix treated as multiple partitions.

**Figure S5.** ASTRAL-III multispecies coalescent phylogeny inferred using the 90% complete matrix.

**Figure S6.** Maximum likelihood phylogeny inferred using the concatenated 90% complete matrix treated as one partition.

**Figure S6.** Maximum likelihood phylogeny inferred using the concatenated 90% complete matrix treated as multiple partitions.

**Figure S7.** Relationships between gene and site concordance factors, log-transformed branch lengths, and ultrafast bootstrap support values estimated using the 75% (top) and 90% (bottom) complete matrices.

**Figure S8. Divergence times and historical biogeography of *Ovalentaria* I: *Incertae Sedis* Lineages.** Time-calibrated phylogeny of *Ovalentaria* from Bayesian node-dating of three sets of 30 UCEs. Bars at nodes indicate 95% highest posterior density intervals of divergence times. Squares at nodes indicate (left) highest probability ancestral area and (right) ancestral habitat estimated in our historical biogeographic reconstruction and polymorphic ancestral state reconstruction. Colors at tips denote tip area and habitat states. Photographs of fishes are by Zachary Randall (Florida Museum of Natural History, USA) and the late John E. Randall.

**Figure S9. Divergence times and historical biogeography of *Ovalentaria* II: *Atheriniformes*.** Time-calibrated phylogeny of *Ovalentaria* from Bayesian node-dating of three sets of 30 UCEs. Bars at nodes indicate 95% highest posterior density intervals of divergence times. Squares at nodes indicate (left) highest probability ancestral area and (right) ancestral habitat estimated in our historical biogeographic reconstruction and polymorphic ancestral state reconstruction. Colors at tips denote tip area and habitat states. Photographs of fishes are by Zachary Randall (Florida Museum of Natural History, USA) and the late John E. Randall.

**Figure S10. Divergence times and historical biogeography of *Ovalentaria* III: *Pholidichthys* and the oldest divergences in *Cichlidae*.** Time-calibrated phylogeny of *Ovalentaria* from Bayesian node-dating of three sets of 30 UCEs. Bars at nodes indicate 95% highest posterior density intervals of divergence times. Squares at nodes indicate (left) highest probability ancestral area and (right) ancestral habitat estimated in our historical biogeographic reconstruction and polymorphic ancestral state reconstruction. Colors at tips denote tip area and habitat states. Photographs of fishes are by Zachary Randall (Florida Museum of Natural History, USA).

**Figure S11. Divergence times and historical biogeography of *Ovalentaria* IV: H-Lineage *Cichlidae*.** Time-calibrated phylogeny of *Ovalentaria* from Bayesian node-dating of three sets of 30 UCEs. Bars at nodes indicate 95% highest posterior density intervals of divergence times. Squares at nodes indicate (left) highest probability ancestral area and (right) ancestral habitat estimated in our historical biogeographic reconstruction and polymorphic ancestral state reconstruction. Colors at tips denote tip area and habitat states. Photographs of fishes are by Zachary Randall (Florida Museum of Natural History, USA).

**Figure S12.** Comparisons of divergence time estimates found in this study and previous phylogenomic analyses. Error bars are 95% highest posterior density intervals, and dots denote median divergence times. Maps of the Earth by Christopher Scotese.

**Figure S13. Salinity Tolerance in *Ovalentaria*.** (A) Distribution of MRCA times (scaled to equal height) across major lineages of *Ovalentaria*. Colors denote clades reconstructed as salinity tolerant (blue) or intolerant of ≥ 30 PSU salinity (orange) in ancestral state reconstructions at right, from both the all-rates-different (ARD) and forward irreversible (i.e., no reversions to being tolerant of ≥ 30 PSU salinity) trait evolution models. (C) Plots show distributions of ages across 20,000 randomly sampled posterior trees from the Bayesian node-dating analysis for clades reconstructed to not tolerate ≥ 30 PSU salinity and the nodes immediately subtending them. Note the long (> 2 Ma) lag times for a loss of salinity tolerance.

**Figure S14. Accumulation of diversity in *Ovalentaria* across space.** (A) Density of dispersals to and from areas, (B) proportion of dispersals to and from areas, and (C) counts of event type per area estimated using biogeographic stochastic mapping.

**Figure S15. Diversification of *Ovalentaria*.** (A) MiSSE diversification phylogram estimated using the best-fitting model (model 6, three hidden states), showing differences in diversification across lineages of *Ovalentaria*, with rapidly diversifying lineages highlighted. (B) Distribution of DR statistics for selected major lineages of *Ovalentaria.*

**Figure S16. Ancestral State Reconstructions.** Full ancestral state reconstructions of tolerance to ≥ 30 PSU salinity estimated under the all-rates-different (ARD, left) and forward irreversible (Irr, right) models. Pie charts at nodes denote probability that each node was ancestrally tolerant (blue) or intolerant (orange) of ≥ 30 PSU salinity.

**Table S1. Specimen Numbers.** Tissues newly extracted for this study are listed below alongside genomes newly skimmed for UCEs. Species not listed are from Ghezelayagh et al. (2022) *Nature Ecology and Evolution*. The two asterisks ** denote a genome that is misidentified on GenBank as a killifish but belongs to the Neotropical cichlid *Apistogramma* sp. YFTC = Yale Fish and Tissue Collection.

| Taxon | Genbank ID or Specimen Voucher # | Barcode Match |
|---|---|---|
| *Salaria pavo* | GCA 043647535.1 | - |
| *Salarias fasciatus* | GCA 902148845.1 | - |
| *Lipophrys pholis* | GCA 963383615.1 | - |
| *Gouania willdenowi* | GCA 900634775.2 | - |
| *Cristiceps australis* | GCA 049082045.1 | - |
| *Blennius ocellaris* | GCA 963422515.1 | - |
| *Lepadogaster candolii* | GCA 965278445.1 | - |
| *Parablennius gattorugine* | GCA 965278395.1 | - |
| *Coryphoblennius galerita* | GCA 965229965.1 | - |
| *Tripterygion tripteronotum* | GCA 049638825.1 | - |
| *Malacoctenus gilli* | GCA 032164315.1 | - |
| *Parablennius parvicornis* | GCA 900302745.1 | - |
| *Malacoctenus tetranemus* | GCA 032274765.1 | - |
| *Aulonocara stuartgranti* | GCA 965226115.1 | - |
| *Oreochromis niloticus* | GCA 001858045.3 | - |
| *Etroplus suratensis* | GCA 041004005.1 | - |
| *Maylandia zebra* | GCA 000238955.5 | - |
| *Parachromis managuensis* | GCA 040437545.1 | - |
| *Astatotilapia calliptera* | GCA 900246225.5 | - |
| *Oreochromis aureus* | GCA 013358895.1 | - |
| *Sarotherodon galilaeus* | GCA 046562855.1 | - |
| *Symphysodon aequifasciata* | GCA 047652445.1 | - |
| *Amphilophus citrinellus* | GCA 013435755.1 | - |
| *Neolamprologus multifasciatus* | GCA 963576455.2 | - |
| *Petenia splendida* | GCA 040437555.1 | - |
| *Pholidichthys leucotaenia* | GCA 020510985.1 | - |

| | | |
|---|---|---|
| *Archocentrus centrarchus* | GCA 007364275.2 | - |
| *Diplotaxodon limnothrissa* | GCA 965183825.1 | - |
| *Labeotropheus trewavasae* | GCA 047663975.1 | - |
| *Chromidotilapia guntheri* | GCA 040805835.1 | - |
| *Pelmatolapia mariae* | GCA 036321145.2 | - |
| *Oreochromis mossambicus* | GCA 922035795.1 | - |
| *Haplochromis burtoni* | GCA 018398535.1 | - |
| *Pundamilia nyererei* | GCA 000239375.1 | - |
| *Mchenga conophoros* | GCA 000150855.1 | - |
| *Neolamprologus brichardi* | GCA 000239395.1 | - |
| *Perissodus microlepis* | GCA 015105195.1 | - |
| *Rhamphochromis esox* | GCA 000150935.1 | - |
| *Pungu maclareni* | GCA 041757325.1 | - |
| *Simochromis diagramma* | GCA 900408965.1 | - |
| *Oreochromis grahami* | GCA 029532605.1 | - |
| *Pterophyllum scalare* | GCA 024447675.2 | - |
| *Haplochromis chilotes* | GCA 018403475.1 | - |
| *Oreochromis spilurus* | GCA 008269305.1 | - |
| *Etroplus canarensis* | GCA 015106775.1 | - |
| *Telmatochromis temporalis* | GCA 015102735.1 | - |
| *Ophthalmotilapia ventralis* | GCA 015105175.1 | - |
| *Labeotropheus fuelleborni* | GCA 000150875.1 | - |
| *Melanochromis auratus* | GCA 000150895.1 | - |
| *Otopharynx argyrosoma* | GCA 046255105.1 | - |
| *Copadichromis chrysonotus* | GCA 050230035.1 | - |
| *Petrochromis trewavasae* | GCA 902810495.1 | - |
| *Tropheus moorii* | GCA 902810505.1 | - |
| *Haplochromis sauvagei* | GCA 018403495.1 | - |
| *Lithochromis rufus* | GCA 018403515.1 | - |
| *Sarotherodon lohbergeri* | GCA 015708405.1 | - |
| *Cichlasoma orientale* | GCA 034780415.1 | - |
| *Pterochromis congicus* | GCA 049180875.1 | - |
| *Oreochromis tanganicae* | GCA 015052225.1 | - |
| *Eretmodus cyanostictus* | GCA 015107895.1 | - |
| *Tilapia sparmanii* | GCA 015108415.1 | - |
| *Hemichromis elongatus* | GCA 015108515.1 | - |
| *Etia nguti* | GCA 015106755.1 | - |
| *Greenwoodochromis staneri* | GCA 015104515.1 | - |
| *Eretmodus marksmithi* | GCA 015110455.1 | - |
| *Haplotaxodon microlepis* | GCA 015105255.1 | - |

| | | |
|---|---|---|
| *Tropheus annectens* | GCA 015106575.1 | - |
| *Coptodon rendalli* | GCA 015108555.1 | - |
| *Perissodus multidentatus* | GCA 015106075.1 | - |
| *Spathodus marlieri* | GCA 015110135.1 | - |
| *Plecodus straeleni* | GCA 015105735.1 | - |
| *Pseudosimochromis babaulti* | GCA 015107015.1 | - |
| *Grammatotria lemairii* | GCA 015107505.1 | - |
| *Greenwoodochromis bellcrossi* | GCA 015106255.1 | - |
| *Petrochromis orthognathus* | GCA 015110415.1 | - |
| *Pseudosimochromis marginatus* | GCA 015052565.1 | - |
| *Serranochromis macrocephalus* | GCA 015106695.1 | - |
| *Boulengerochromis microlepis* | GCA 015107555.1 | - |
| *Coptodon congicus* | GCA 049180755.1 | - |
| *Perissodus eccentricus* | GCA 015107955.1 | - |
| *Neolamprologus furcifer* | GCA 015108955.1 | - |
| *Cyprichromis coloratus* | GCA 015107215.1 | - |
| *Neolamprologus longior* | GCA 015109375.1 | - |
| *Tanganicodus irsacae* | GCA 015110155.1 | - |
| *Lepidiolamprologus cunningtoni* | GCA 015105495.1 | - |
| *Neolamprologus variostigma* | GCA 015108695.1 | - |
| *Variabilichromis moorii* | GCA 015102225.1 | - |
| *Sargochromis carlottae* | GCA 015106635.1 | - |
| *Thoracochromis brauschi* | GCA 015106615.1 | - |
| *Neolamprologus helianthus* | GCA 015108395.1 | - |
| *Paracyprichromis nigripinnis* | GCA 015103275.1 | - |
| *Tropheus polli* | GCA 015111015.1 | - |
| *Pelvicachromis taeniatus* | GCA 015108175.1 | - |
| *Neolamprologus toae* | GCA 015110075.1 | - |
| *Sarotherodon steinbachi* | GCA 015708585.1 | - |
| *Coptodon bakossiorum* | GCA 015106655.1 | - |
| *Xenotilapia flavipinnis* | GCA 015105975.1 | - |
| *Neolamprologus gracilis* | GCA 015108275.1 | - |
| *Neolamprologus ventralis* | GCA 015109735.1 | - |
| *Greenwoodochromis christyi* | GCA 015107935.1 | - |
| *Cyprichromis zonatus* | GCA 015052445.1 | - |
| *Pseudosimochromis curvifrons* | GCA 015052425.1 | - |
| *Xenotilapia nigrolabiata* | GCA 015106095.1 | - |
| *Australoheros facetus* | GCA 030220045.1 | - |
| *Neolamprologus cylindricus* | GCA 015103235.1 | - |
| *Orthochromis indermauri* | GCA 015103075.1 | - |

| | | |
|---|---|---|
| *Neolamprologus caudopunctatus* | GCA 015105295.1 | - |
| *Hemibates koningsi* | GCA 015052305.1 | - |
| *Xenochromis hecqui* | GCA 015107655.1 | - |
| *Lepidiolamprologus kendalli* | GCA 015102695.1 | - |
| *Neolamprologus tetracanthus* | GCA 015105335.1 | - |
| *Xenotilapia longispinis* | GCA 015109775.1 | - |
| *Spathodus erythrodon* | GCA 015108895.1 | - |
| *Tangachromis dhanisi* | GCA 015113095.1 | - |
| *Julidochromis marlieri* | GCA 015108065.1 | - |
| *Neolamprologus devosi* | GCA 015052545.1 | - |
| *Neolamprologus kungweensis* | GCA 015110375.1 | - |
| *Ectodus descampsii* | GCA 015104995.1 | - |
| *Baileychromis centropomoides* | GCA 015107735.1 | - |
| *Neolamprologus modestus* | GCA 015102555.1 | - |
| *Pharyngochromis acuticeps* | GCA 015108475.1 | - |
| *Julidochromis dickfeldi* | GCA 015105015.1 | - |
| *Lepidiolamprologus pleuromaculatus* | GCA 015109975.1 | - |
| *Cyphotilapia gibberosa* | GCA 015105655.1 | - |
| *Xenotilapia rotundiventralis* | GCA 015105935.1 | - |
| *Cardiopharynx schoutedeni* | GCA 015109795.1 | - |
| *Paratilapia polleni* | GCA 015106735.1 | - |
| *Petrochromis horii* | GCA 015106495.1 | - |
| *Neolamprologus longicaudatus* | GCA 015108155.1 | - |
| *Lamprologus callipterus* | GCA 015052345.1 | - |
| *Lamprologus speciosus* | GCA 015109675.1 | - |
| *Neolamprologus nigriventris* | GCA 015102355.1 | - |
| *Reganochromis calliurus* | GCA 015104255.1 | - |
| *Neolamprologus falcicula* | GCA 015110515.1 | - |
| *Julidochromis transcriptus* | GCA 015103115.1 | - |
| *Astatotilapia stappersii* | GCA 015110235.1 | - |
| *Neolamprologus mondabu* | GCA 015106975.1 | - |
| *Lobochilotes labiata* | GCA 015104595.1 | - |
| *Telmatochromis vittatus* | GCA 015107575.1 | - |
| *Greenwoodochromis abeelei* | GCA 015052385.1 | - |
| *Lamprologus ornatipinnis* | GCA 015109935.1 | - |
| *Bathybates hornii* | GCA 015107495.1 | - |
| *Stomatepia pindu* | GCA 015708495.1 | - |
| *Neolamprologus pectoralis* | GCA 015108755.1 | - |
| *Lamprologus signatus* | GCA 015106455.1 | - |
| *Neolamprologus meeli* | GCA 015107355.1 | - |

| | | |
|---|---|---|
| *Bujurquina vittata* | GCA 015106715.1 | - |
| *Neolamprologus olivaceous* | GCA 015108255.1 | - |
| *Plecodus paradoxus* | GCA 015107295.1 | - |
| *Heterotilapia buettikoferi* | GCA 015108575.1 | - |
| *Neolamprologus walteri* | GCA 015109215.1 | - |
| *Xenotilapia caudafasciata* | GCA 015106215.1 | - |
| *Neolamprologus leleupi* | GCA 015108215.1 | - |
| *Julidochromis marksmithi* | GCA 015110755.1 | - |
| *Lamprologus tigripictilis* | GCA 015108455.1 | - |
| *Lamprologus laparogramma* | GCA 015107315.1 | - |
| *Heterochromis multidens* | GCA 015108495.1 | - |
| *Neolamprologus marunguensis* | GCA 015108335.1 | - |
| *Callochromis pleurospilus* | GCA 015110035.1 | - |
| *Shuja horei* | GCA 015103775.1 | - |
| *Interochromis loocki* | GCA 015105475.1 | - |
| *Neolamprologus pulcher* | GCA 015104715.1 | - |
| *Neolamprologus hecqui* | GCA 015102335.1 | - |
| *Xenotilapia leptura* | GCA 015105755.1 | - |
| *Petrochromis macrognathus* | GCA 015111435.1 | - |
| *Tylochromis polylepis* | GCA 015107715.1 | - |
| *Neolamprologus savoryi* | GCA 015104745.1 | - |
| *Xenotilapia melanogenys* | GCA 015103415.1 | - |
| *Xenotilapia tenuidentata* | GCA 015118035.1 | - |
| *Lepidiolamprologus profundicola* | GCA 015109915.1 | - |
| *Sarotherodon caroli* | GCA 015708515.1 | - |
| *Altolamprologus compressiceps* | GCA 015104655.1 | - |
| *Paracyprichromis brieni* | GCA 015110335.1 | - |
| *Gnathochromis permaxillaris* | GCA 015104535.1 | - |
| *Benthochromis horii* | GCA 015102815.1 | - |
| *Neolamprologus crassus* | GCA 015104075.1 | - |
| *Neolamprologus fasciatus* | GCA 015102195.1 | - |
| *Bathybates fasciatus* | GCA 015104405.1 | - |
| *Orthochromis malagaraziensis* | GCA 015111915.1 | - |
| *Xenotilapia papilio* | GCA 015106585.1 | - |
| *Tropheus brichardi* | GCA 015110215.1 | - |
| *Telmatochromis bifrenatus* | GCA 015111975.1 | - |
| *Neolamprologus bifasciatus* | GCA 015113255.1 | - |
| *Neolamprologus buescheri* | GCA 015107095.1 | - |
| *Orthochromis uvinzae* | GCA 015111895.1 | - |
| *Neolamprologus chitamwebwai* | GCA 015109035.1 | - |

| | | |
|---|---|---|
| *Cyphotilapia frontosa* | GCA 015110975.1 | - |
| *Benthochromis melanoides* | GCA 015102775.1 | - |
| *Altolamprologus calvus* | GCA 015052405.1 | - |
| *Xenotilapia nasus* | GCA 015102635.1 | - |
| *Lamprologus similis* | GCA 015109475.1 | - |
| *Neolamprologus calliurus* | GCA 015103035.1 | - |
| *Xenotilapia ornatipinnis* | GCA 015110015.1 | - |
| *Xenotilapia bathyphilus* | GCA 015104175.1 | - |
| *Xenotilapia boulengeri* | GCA 015105375.1 | - |
| *Xenotilapia sima* | GCA 015102085.1 | - |
| *Orthochromis mazimeroensis* | GCA 015109575.1 | - |
| *Astatoreochromis straeleni* | GCA 015109875.1 | - |
| *Neolamprologus obscurus* | GCA 015102675.1 | - |
| *Cyprichromis leptosoma* | GCA 015104635.1 | - |
| *Tropheus duboisi* | GCA 015109155.1 | - |
| *Xenotilapia ochrogenys* | GCA 015106855.1 | - |
| *Limnochromis auritus* | GCA 015104565.1 | - |
| *Neolamprologus christyi* | GCA 015104465.1 | - |
| *Telmatochromis brachygnathus* | GCA 015107635.1 | - |
| *Callochromis macrops* | GCA 015103395.1 | - |
| *Chalinochromis cyanophleps* | GCA 015112635.1 | - |
| *Bathybates graueri* | GCA 015102795.1 | - |
| *Neolamprologus leloupi* | GCA 015111655.1 | - |
| *Apistogramma diplotaenia* | GCA 015052665.1 | - |
| *Cyprichromis microlepidotus* | GCA 015106915.1 | - |
| *Trematochromis benthicola* | GCA 015103355.1 | - |
| *Cyathopharynx furcifer* | GCA 015103655.1 | - |
| *Neolamprologus petricola* | GCA 015112555.1 | - |
| *Neolamprologus niger* | GCA 015111955.1 | - |
| *Oreochromis malagarasi* | GCA 015111835.1 | - |
| *Triglachromis otostigma* | GCA 015107115.1 | - |
| *Lepidiolamprologus kamambae* | GCA 015110795.1 | - |
| *Cyprichromis pavo* | GCA 015052265.1 | - |
| *Plecodus elaviae* | GCA 015113155.1 | - |
| *Neolamprologus mustax* | GCA 015102955.1 | - |
| *Julidochromis ornatus* | GCA 015104675.1 | - |
| *Trematocara macrostoma* | GCA 015104355.1 | - |
| *Lestradea stappersii* | GCA 015106895.1 | - |
| *Lepidiolamprologus attenuatus* | GCA 015103955.1 | - |
| *Gnathochromis pfefferi* | GCA 015052505.1 | - |

| | | |
|---|---|---|
| *Ophthalmotilapia nasuta* | GCA 015103715.1 | - |
| *Bathybates minor* | GCA 015107595.1 | - |
| *Hemibates stenosoma* | GCA 015106175.1 | - |
| *Chalinochromis brichardi* | GCA 015102055.1 | - |
| *Cyathopharynx foae* | GCA 015105635.1 | - |
| *Telmatochromis dhonti* | GCA 015111745.1 | - |
| *Telmatochromis brichardi* | GCA 015108855.1 | - |
| *Chalinochromis popelini* | GCA 015103555.1 | - |
| *Trematocara unimaculatum* | GCA 015106295.1 | - |
| *Xenotilapia spiloptera* | GCA 015103795.1 | - |
| *Neolamprologus boulengeri* | GCA 015109695.1 | - |
| *Haplotaxodon trifasciatus* | GCA 015105275.1 | - |
| *Haplochromis vanheusdeni* | GCA 015106555.1 | - |
| *Trematocara nigrifrons* | GCA 015104335.1 | - |
| *Limnotilapia dardennii* | GCA 015103815.1 | - |
| *Petrochromis polyodon* | GCA 015103915.1 | - |
| *Apistogramma caetei* | GCA 050511875.1 | - |
| *Trematocara marginatum* | GCA 015104795.1 | - |
| *Benitochromis conjunctus* | GCA 015052645.1 | - |
| *Aulonocranus dewindti* | GCA 015102295.1 | - |
| *Trematocara stigmaticum* | GCA 015107995.1 | - |
| *Petrochromis famula* | GCA 015108095.1 | - |
| *Neolamprologus timidus* | GCA 015112655.1 | - |
| *Petrochromis fasciolatus* | GCA 015103155.1 | - |
| *Bathybates vittatus* | GCA 015107455.1 | - |
| *Neolamprologus tretocephalus* | GCA 015109125.1 | - |
| *Ophthalmotilapia heterodonta* | GCA 015103535.1 | - |
| *Neolamprologus brevis* | GCA 015103015.1 | - |
| *Haplochromis paludinosus* | GCA 015111855.1 | - |
| *Lestradea perspicax* | GCA 015105095.1 | - |
| *Bathybates ferox* | GCA 015111595.1 | - |
| *Bathybates leo* | GCA 015102875.1 | - |
| *Lepidiolamprologus elongatus* | GCA 015102115.1 | - |
| *Julidochromis regani* | GCA 015109195.1 | - |
| *Andinoacara biseriatus* | GCA 015108635.1 | - |
| *Lamprologus stappersi* | GCA 015108715.1 | - |
| *Neolamprologus sexfasciatus* | GCA 015102515.1 | - |
| *Ophthalmotilapia boops* | GCA 015110735.1 | - |
| *Lepidiolamprologus mimicus* | GCA 015111395.1 | - |
| *Neolamprologus prochilus* | GCA 015106535.1 | - |

| | | |
|---|---|---|
| *Lamprologus ocellatus* | GCA 015052485.1 | - |
| *Callochromis melanostigma* | GCA 015109845.1 | - |
| *Benthochromis tricoti* | GCA 015111415.1 | - |
| *Pseudocrenilabrus philander* | GCA 032270725.1 | - |
| *Astatotilapia flaviijosephi* | GCA 015112995.1 | - |
| *Lamprologus lemairii* | GCA 015107875.1 | - |
| *Trematocara zebra* | GCA 015110815.1 | - |
| *Cunningtonia longiventralis* | GCA 015106375.1 | - |
| *Xenotilapia singularis* | GCA 015104975.1 | - |
| *Neolamprologus cancellatus* | GCA 015113075.1 | - |
| *Australoheros scitulus* | GCA 015108615.1 | - |
| *Amphilophus zaliosus* | GCA 015108585.1 | - |
| *Trematocara variabile* | GCA 015103455.1 | - |
| *Ptychochromis oligacanthus* | GCA 015103295.1 | - |
| *Trematocara caparti* | GCA 015112415.1 | - |
| *Parambassis ranga* | GCA 900634625.2 | - |
| *Ambassis kopsii* | GCA 050573535.1 | - |
| *Ambassis urotaenia* | GCA 049180735.1 | - |
| *Ambassis buruensis* | GCA 050573475.1 | - |
| *Ambassis agassizii* | GCA 036025115.1 | - |
| *Mugil cephalus* | GCA 022458985.1 | - |
| *Chelon labrosus* | GCA 963514085.1 | - |
| *Planiliza haematocheilus* | GCA 005024645.1 | - |
| *Ellochelon vaigiensis* | GCA 041107635.1 | - |
| *Chelon auratus* | GCA 030220025.1 | - |
| *Pseudochromis fuscus* | GCA 900323345.1 | - |
| *Amphiprion ocellaris* | GCA 022539595.1 | - |
| *Acanthochromis polyacanthus* | GCA 021347895.1 | - |
| *Amphiprion percula* | GCA 003047355.2 | - |
| *Amphiprion clarkii* | GCA 027123335.1 | - |
| *Dascyllus trimaculatus* | GCA 024666655.1 | - |
| *Stegastes partitus* | GCA 000690725.1 | - |
| *Chrysiptera cyanea* | GCA 046866775.1 | - |
| *Chromis chromis* | GCA 900302755.1 | - |
| *Abudefduf troschelii* | GCA 043656515.1 | - |
| *Dascyllus aruanus* | GCA 049176935.1 | - |
| *Pomacentrus brachialis* | GCA 049176895.1 | - |
| *Embiotoca jacksoni* | GCA 022577435.1 | - |
| *Brachyistius frenatus* | GCA 029030835.1 | - |
| *Oryzias latipes* | GCA 002234675.1 | - |

| | | |
|---|---|---|
| *Nothobranchius furzeri* | GCA 043380555.1 | - |
| *Fundulus heteroclitus* | GCA 011125445.2 | - |
| *Oryzias melastigma* | GCA 002922805.2 | - |
| *Poecilia reticulata* | GCA 000633615.2 | - |
| *Kryptolebias marmoratus* | GCA 001649575.2 | - |
| *Gambusia affinis* | GCA 019740435.1 | - |
| *Gambusia holbrooki* | GCA 965282415.1 | - |
| *Xiphophorus maculatus* | GCA 002775205.2 | - |
| *Cololabis saira* | GCA 033807715.1 | - |
| *Menidia menidia* | GCA 048628825.1 | - |
| *Oryzias javanicus* | GCA 003999625.1 | - |
| *Oryzias curvinotus* | GCA 023969325.1 | - |
| *Oryzias sinensis* | GCA 037389245.1 | - |
| *Xiphophorus cortezi* | GCA 041380905.1 | - |
| *Xiphophorus birchmanni* | GCA 036418095.1 | - |
| *Anableps anableps* | GCA 014839685.1 | - |
| *Xiphophorus couchianus* | GCA 001444195.3 | - |
| *Odontesthes bonariensis* | GCA 027942865.1 | - |
| *Xiphophorus hellerii* | GCA 003331165.2 | - |
| *Nematolebias whitei* | GCA 014905685.2 | - |
| *Cyprinodon nevadensis mionectes* | GCA 030533455.1 | - |
| *Neostethus bicornis* | GCA 902685375.1 | - |
| *Xiphophorus malinche* | GCA 041429825.1 | - |
| *Girardinichthys multiradiatus* | GCA 021462225.2 | - |
| *Cyprinodon diabolis* | GCA 030533445.1 | - |
| *Fundulus diaphanus* | GCA 037039145.1 | - |
| *Melanotaenia boesemani* | GCA 017639745.1 | - |
| *Poecilia picta* | GCA 033032245.1 | - |
| *Xenentodon cancila* | GCA 014839995.1 | - |
| *Telmatherina bonti* | GCA 933228915.1 | - |
| *Kryptolebias hermaphroditus* | GCA 007896545.1 | - |
| *Valencia hispanica* | GCA 963556495.2 | - |
| *Atherina presbyter* | GCA 965295175.1 | - |
| *Hyporhamphus sajori* | GCA 050306495.1 | - |
| *Hyporhamphus intermedius* | GCA 050306515.1 | - |
| *Melanotaenia duboulayi* | GCA 026261665.1 | - |
| *Fundulus grandis* | GCA 049004275.1 | - |
| *Poecilia formosa* | GCA 000485575.1 | - |
| *Poecilia mexicana* | GCA 001443325.1 | - |
| *Poecilia latipinna* | GCA 001443285.1 | - |

| | | |
|---|---|---|
| *Leuresthes tenuis* | GCA 036924035.1 | - |
| *Cyprinodon variegatus* | GCA 000732505.1 | - |
| *Austrofundulus limnaeus* | GCA 001266775.1 | - |
| *Poeciliopsis prolifica* | GCA 027474105.1 | - |
| *Menidia beryllina* | GCA 013363375.1 | - |
| *Chirostoma estor* | GCA 043091105.1 | - |
| *Nothobranchius kuhntae* | GCA 006942095.1 | - |
| *Melanotaenia fluviatilis* | GCA 036025615.1 | - |
| *Melanotaenia splendida* | GCA 037177455.1 | - |
| *Oryzias celebensis* | GCA 014656515.1 | - |
| *Fundulus olivaceus* | GCA 049004345.1 | - |
| *Poeciliopsis turneri* | GCA 922788865.1 | - |
| *Cyprinodon tularosa* | GCA 016077235.1 | - |
| *Atherinomorus stipes* | GCA 023101725.1 | - |
| *Cyprinodon brontotheroides* | GCA 018398635.1 | - |
| *Poeciliopsis presidionis* | GCA 903068095.1 | - |
| *Austrolebias charrua* | GCA 038048925.1 | - |
| *Atherinosoma microstoma* | GCA 037177055.1 | - |
| *Melanotaenia parkinsoni* | GCA 036919975.1 | - |
| *Craterocephalus eyresii* | GCA 037177035.1 | - |
| *Melanotaenia australis* | GCA 037177175.1 | - |
| *Craterocephalus fluviatilis* | GCA 037177075.1 | - |
| *Apricaphanius iberus* | GCA 028564705.1 | - |
| *Oryzias mekongensis* | GCA 050043955.1 | - |
| *Poecilia sulphuraria* | GCA 051168745.1 | - |
| *Poeciliopsis retropinna* | GCA 010277075.1 | - |
| *Phalloptychus januarius* | GCA 902982915.1 | - |
| *Oryzias luzonensis* | GCA 050043775.1 | - |
| *Poeciliopsis gracilis* | GCA 911649635.1 | - |
| *Hirundichthys speculiger* | GCA 018398805.1 | - |
| *Fundulus xenicus* | GCA 049004235.1 | - |
| *Fundulus catenatus* | GCA 049004365.1 | - |
| *Crenichthys baileyi* | GCA 036173005.1 | - |
| *Aphyosemion australe* | GCA 006937985.1 | - |
| *Poeciliopsis turrubarensis* | GCA 010277015.1 | - |
| *Adrianichthys oophorus* | GCA 036245035.1 | - |
| *Pachypanchax playfairii* | GCA 006937955.1 | - |
| *Poeciliopsis paucimaculata* | GCA 903067055.1 | - |
| *Phalloptychus iheringii* | GCA 046118055.1 | - |
| *Poeciliopsis infans* | GCA 903067105.1 | - |

| | | |
|---|---|---|
| *Poeciliopsis occidentalis* | GCA 010883035.1 | - |
| *Alfaro cultratus* | GCA 903066825.1 | - |
| *Callopanchax toddi* | GCA 006937965.1 | - |
| *Girardinus metallicus* | GCA 903067095.1 | - |
| *Brachyrhaphis roseni* | GCA 903066815.1 | - |
| *Phalloceros buckupi* | GCA 046118275.1 | - |
| *Phalloceros spiloura* | GCA 046118235.1 | - |
| *Cnesterodon brevirostratus* | GCA 046269305.1 | - |
| *Orestias gloriae* | GCA 032876255.1 | - |
| *Orestias chungarensis* | GCA 032876305.1 | - |
| *Orestias laucaensis* | GCA 032876265.1 | - |
| *Oryzias dopingdopingensis* | GCA 030512385.1 | - |
| *Poecilia gillii* | GCA 903068115.1 | - |
| *Phalloceros leptokeras* | GCA 046118015.1 | - |
| *Micropoecilia bifurca* | GCA 903067075.1 | - |
| *Xenoophorus captivus* | GCA 040091335.1 | - |
| *Pamphorichthys hollandi* | GCA 046118195.1 | - |
| *Phalloceros titthos* | GCA 046118035.1 | - |
| *Characodon lateralis* | GCA 036250325.1 | - |
| *Kryptolebias ocellatus* | GCA 028566035.1 | - |
| *Ameca splendens* | GCA 040091285.1 | - |
| *Poecilia velifera* | GCA 036873795.1 | - |
| *Poecilia wingei* | GCA 903068105.1 | - |
| *Ataeniobius toweri* | GCA 036250305.1 | - |
| *Kryptolebias gracilis* | GCA 028565975.1 | - |
| *Goodea atripinnis* | GCA 040091325.1 | - |
| *Gambusia yucatana* | GCA 035584275.1 | - |
| *Xenotaenia resolanae* | GCA 040091345.1 | - |
| *Phalloceros megapolos* | GCA 046118095.1 | - |
| *Arrhamphus sclerolepis* | GCA 037176975.1 | - |
| *Zenarchopterus dispar* | GCA 028564865.1 | - |
| *Phalloceros reisi* | GCA 046118115.1 | - |
| *Rhadinocentrus ornatus* | GCA 037177495.1 | - |
| *Phalloceros maldonadoi* | GCA 046118075.1 | - |
| *Melanotaenia trifasciata* | GCA 037177355.1 | - |
| *Jordanella pulchra* | GCA 034780135.1 | - |
| *Ilyodon furcidens* | GCA 040091305.1 | - |
| *Melanotaenia pygmaea* | GCA 036919955.1 | - |
| *Cynopoecilus melanotaenia* | GCA 040207505.1 | - |
| *Cynodonichthys magdalenae* | GCA 031762555.1 | - |

| | | |
|---|---|---|
| *Anablepsoides urophthalmus* | GCA 050573655.1 | - |
| *Melanorivulus punctatus* | GCA 051253435.1 | - |
| *Craterocephalus amniculus* | GCA 036024015.2 | - |
| *Pseudomugil tenellus* | GCA 037177515.1 | - |
| *Melanotaenia maccullochi* | GCA 037177255.1 | - |
| *Melanotaenia nigrans* | GCA 037177435.1 | - |
| *Leptatherina wallacei* | GCA 028567055.1 | - |
| *Melanorivulus apiamici* | GCA 050511755.1 | - |
| *Kiunga ballochi* | GCA 037177335.1 | - |
| *Melanorivulus atlanticus* | GCA 044758275.1 | - |
| *Pseudomugil gertrudae* | GCA 037177535.1 | - |
| *Melanorivulus parnaibensis* | GCA 043658355.1 | - |
| *Melanotaenia catherinae* | GCA 037177155.1 | - |
| *Kryptolebias brasiliensis* | GCA 028566095.1 | - |
| *Pituna xinguensis* | GCA 051253915.1 | - |
| *Atherinomorus endrachtensis* | GCA 050573555.1 | - |
| *Melanotaenia monticola* | GCA 036919915.1 | - |
| *Brachyrhaphis terrabensis* | GCA 903067065.1 | - |
| *Doboatherina balabacensis* | GCA 049181575.1 | - |
| *Simpsonichthys santanae* | GCA 031762415.1 | - |
| *Anablepsoides cearensis* | GCA 032275365.1 | - |
| *Melanorivulus jalapensis* | GCA 044474325.1 | - |
| *Nematolebias papilliferus* | GCA 031762515.1 | - |
| *Millerichthys robustus* | GCA 034783495.1 | - |
| *Atlantirivulus luelingi* | GCA 051252835.1 | - |
| *Apistogramma* sp.** | GCA 050511815.1 | - |
| *Hypsolebias longignatus* | GCA 031762435.1 | - |
| *Cynopoecilus notabilis* | GCA 051253235.1 | - |
| *Anablepsoides waimacui* | GCA 051016645.1 | - |
| *Hypsolebias antenori* | GCA 031762315.1 | - |
| *Ophthalmolebias ilheusensis* | GCA 043658435.1 | - |
| *Cynopoecilus intimus* | GCA 051253015.1 | - |
| *Hypsolebias martinsi* | GCA 031762355.1 | - |
| *Notholebias vermiculatus* | GCA 050511835.1 | - |
| *Atlantirivulus santensis* | GCA 043658295.1 | - |
| *Melanotaenia mubiensis* | GCA 028567035.1 | - |
| *Melanorivulus giarettai* | GCA 051253415.1 | - |
| *Anablepsoides ornatus* | GCA 050511675.1 | - |
| *Atlantirivulus riograndensis* | GCA 051252915.1 | - |
| *Atlantirivulus haraldsiolii* | GCA 050573675.1 | - |

| | | |
|---|---|---|
| *Hypsolebias faouri* | GCA 050511735.1 | - |
| *Hypsolebias coamazonicus* | GCA 032273985.1 | - |
| *Neofundulus paraguayensis* | GCA 051253475.1 | - |
| *Leptolebias minimus* | GCA 051253995.1 | - |
| *Melanotaenia affinis* | GCA 036919935.1 | - |
| *Atlantirivulus jurubatibensis* | GCA 044758845.1 | - |
| *Hypoatherina temminckii* | GCA 051016685.1 | - |
| *Hypsolebias janaubensis* | GCA 050511795.1 | - |
| *Hypsolebias ghisolfii* | GCA 051253275.1 | - |
| *Craterocephalus stercusmuscarum* | GCA 037177115.1 | - |
| *Cynolebias leptocephalus* | GCA 051253775.1 | - |
| *Laimosemion strigatum* | GCA 051254375.1 | - |
| *Melanotaenia caerulea* | GCA 037177135.1 | - |
| *Hypsolebias gongobira* | GCA 044474665.1 | - |
| *Hypsolebias lulai* | GCA 044474745.1 | - |
| *Hypsolebias bonita* | GCA 044758785.1 | - |
| *Hypsolebias shibattai* | GCA 051253735.1 | - |
| *Atherinomorus duodecimalis* | GCA 050573715.1 | - |
| *Melanotaenia eachamensis* | GCA 037177195.1 | - |
| *Leptopanchax itanhaensis* | GCA 050511855.1 | - |
| *Cynodonichthys tenuis* | GCA 031001815.1 | - |
| *Anablepsoides vieirai* | GCA 051253595.1 | - |
| *Argolebias nigripinnis* | GCA 031763805.1 | - |
| *Anablepsoides rubrolineatus* | GCA 031762495.1 | - |
| *Hypsolebias flavicaudatus* | GCA 051253315.1 | - |
| *Atherina boyeri* | GCA 029874935.1 | - |
| *Leptolebias citrinipinnis* | GCA 051253375.1 | - |
| *Craterocephalus marjoriae* | GCA 037177095.1 | - |
| *Cynopoecilus nigrovittatus* | GCA 051253155.1 | - |
| *Atlantirivulus guanabarensis* | GCA 044758625.1 | - |
| *Cynopoecilus fulgens* | GCA 051253175.1 | - |
| *Cynolebias porosus* | GCA 051253795.1 | - |
| *Austrolebias alexandri* | GCA 051253135.1 | - |
| *Austrolebias periodicus* | GCA 051252955.1 | - |
| *Melanotaenia goldiei* | GCA 041107655.1 | - |
| *Atlantirivulus lazzarotoi* | GCA 044758765.1 | - |
| *Atlantirivulus ribeirensis* | GCA 051252935.1 | - |
| *Austrolebias minuano* | GCA 051253195.1 | - |
| *Austrolebias wolterstorffi* | GCA 051252895.1 | - |
| *Austrolebias adloffi* | GCA 051016705.1 | - |

| | | |
|---|---|---|
| *Austrolebias litzi* | GCA 050573775.1 | - |
| *Pseudomugil signifer* | GCA 037177565.1 | - |
| *Melanotaenia vanheurni* | GCA 037177475.1 | - |
| *Melanotaenia kokasensis* | GCA 037177375.1 | - |
| *Assessor macneilli* | YFTC 23731 | Yes |
| *Notograptus guttatus* | YFTC 23728 | Yes |
| *Paraplesiops alisonae* | YFTC 24284 | Yes |
| *Blennodesmus scapularis* | YFTC 23735 | No |
| *Halidesmus scapularis* | YFTC 21677 | Yes |
| *Natalichthys sam* | YFTC 21674 | Yes |
| *Congrogadus subducens* | LEOT 505 | Yes |
| *Congrogadus winterbottomi* | YFTC 23726 | No |
| *Assiculus punctatus* | YFTC 23729 | No |
| *Cypho purpurascens* | YFTC 23732 | Yes |
| *Cypho* sp. | YFTC 21672 | No |
| *Labracinus lineatus* | YFTC 23730 | No |
| *Pseudoplesiops revellei* | YFTC 21676 | Yes |

**Table S2. Model Fitting for Ancestral State reconstructions.** Best-fit model for each analysis is bolded. ER = Equal Rates; ARD = All Rates Different; Irr1 = Forward Irreversible Model, i.e., no regains, Irr2 = Backward Irreversible model, i.e., no secondary losses.

| Fused Pharyngeal Jaw | log(L) | d.f. | AIC | weight |
|---|---|---|---|---|
| ER model | -20.15864 | 1 | 42.31728 | 0.14157588 |
| ARD model | -18.67056 | 2 | 41.34112 | 0.23065319 |
| **Irr1 model** | **-18.67056** | **1** | **39.34112** | **0.62698038** |
| Irr2 model | -25.3465 | 1 | 52.693 | 0.00079055 |

| Freshwater Component | log(L) | d.f. | AIC | weight |
|---|---|---|---|---|
| ER model | -58.89563 | 1 | 119.7913 | 1.99E-01 |
| ARD model | -57.81803 | 2 | 119.6361 | 2.15E-01 |
| Irr1 model | -67.61901 | 1 | 137.238 | 3.24E-05 |
| **Irr2 model** | **-57.81803** | **1** | **117.6361** | **5.85E-01** |

| Marine Component | log(L) | d.f. | AIC | weight |
|---|---|---|---|---|
| **ER model** | **-78.23924** | **1** | **158.4785** | **6.90E-01** |
| ARD model | -78.27746 | 2 | 160.5549 | 2.44E-01 |
| Irr1 model | -80.60019 | 1 | 163.2004 | 6.51E-02 |
| Irr2 model | -110.90875 | 1 | 223.8175 | 4.48E-15 |

| Salinity Tolerance | log(L) | d.f. | AIC | weight |
|---|---|---|---|---|
| ER model | -85.8754 | 1 | 173.7508 | 1.37E-05 |
| **ARD model** | **-73.67688** | **2** | **151.3538** | **1.00E+00** |
| Irr1 model | -96.69615 | 1 | 195.3923 | 2.74E-10 |
| Irr2 model | -87.94029 | 1 | 177.8806 | 1.74E-06 |

**Table S3. Phylogenetic Least Squares of Discrete Traits and DR Statistics.**

| Analysis | Value | Std. Error | t-value | p-value |
|---|---|---|---|---|
| (Intercept) | -3.1501659 | 1.9394352 | -1.6242697 | 0.104887 |
| Freshwater Component | 0.07667037 | 0.2753521 | 0.2784449 | 0.7807748 |
| | Value | Std. Error | t-value | p-value |
| (Intercept) | -3.0886502 | 1.9529149 | -1.58155902 | 0.114321 |
| Fused Pharyngeal Jaw (FPC) | -0.0774095 | 0.8404297 | -0.09210704 | 0.9266463 |
| | Value | Std. Error | t-value | p-value |
| (Intercept) | -3.1272739 | 1.9366068 | -1.6148213 | 0.1069188 |
| FPC + Freshwater Component | 0.1006266 | 0.5701088 | 0.1765043 | 0.8599623 |
| | Value | Std. Error | t-value | p-value |
| (Intercept) | -3.0324528 | 1.2537751 | -2.4186576 | 0.01670717 |
| Tolerance | -0.0694369 | 0.1004817 | -0.6910398 | 0.49054819 |

**Table S4. Hidden State Speciation and Extinction Model Fitting.** Bolded indicates best fit model for each variable examined. Values are AIC scores. Dull Null = model wherein traits do not associate with different diversification rates, BiSSE = binary SSE model, HiSSE CD, character-state dependent hidden state SSE model, HiSSE CID, character-independent hidden state SSE model.

| Variable | Dull Null | BiSSE | HiSSE CD | HiSSE CID |
|---|---|---|---|---|
| Freshwater Component | 3179.794 | 3120.011 | **2930.362** | 2931.904 |
| Fused Pharyngeal Jaw | 3038.35 | 3019.666 | **2780.38** | 2832.614 |
| FPC + Freshwater Component | 3109.37 | 3000.603 | **2786.427** | 2861.482 |
| Tolerance | 1376.545 | 1378.917 | 1388.209 | **1377.408** |

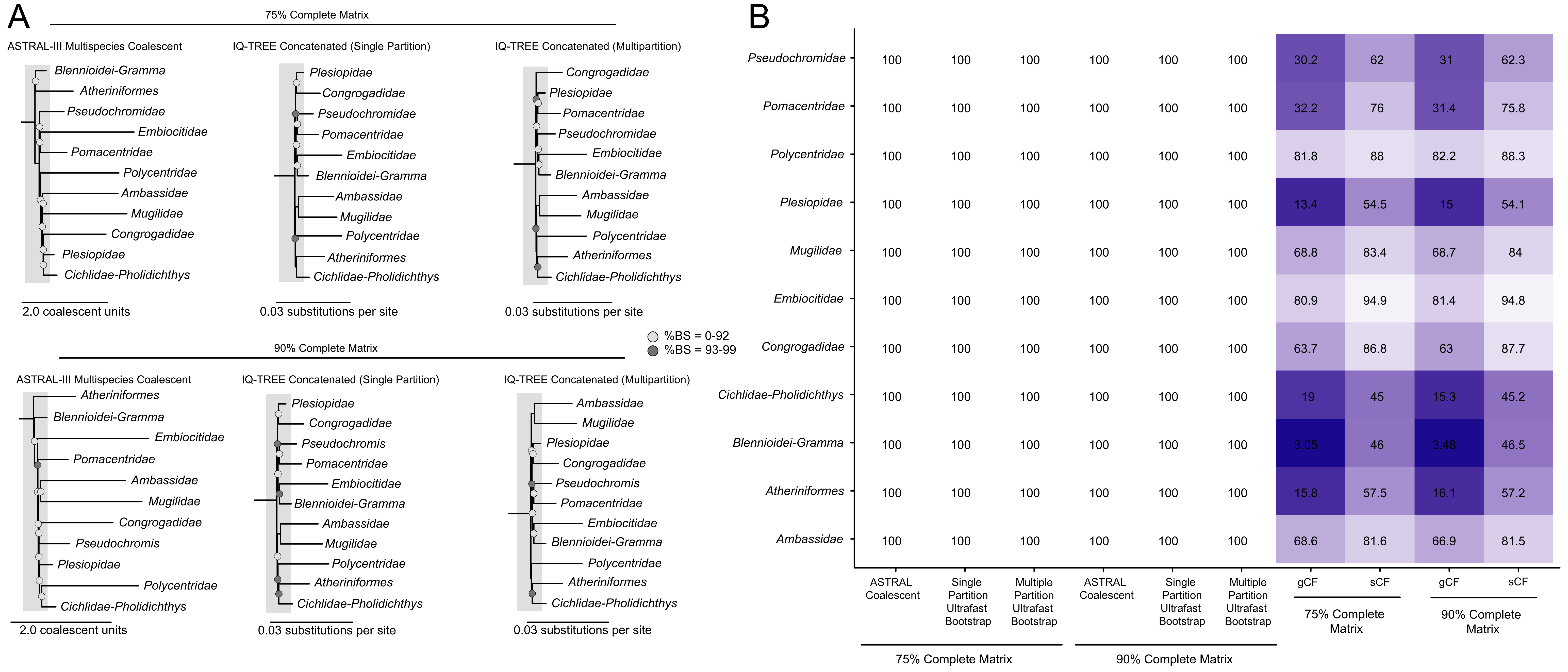

A
75% Complete Matrix
ASTRAL-III Multispecies Coalescent
Blennioidei-Gramma
Atheriniformes
Pseudochromidae
Embiocitidae
Pomacentridae
Polycentridae
Ambassidae
Mugilidae
Congrogadidae
Plesiopidae
Cichlidae-Pholidichthys
2.0 coalescent units
IQ-TREE Concatenated (Single Partition)
Plesiopidae
Congrogadidae
Pseudochromidae
Pomacentridae
Embiocitidae
Blennioidei-Gramma
Ambassidae
Mugilidae
Polycentridae
Atheriniformes
Cichlidae-Pholidichthys
0.03 substitutions per site
IQ-TREE Concatenated (Multipartition)
Congrogadidae
Plesiopidae
Pomacentridae
Pseudochromidae
Embiocitidae
Blennioidei-Gramma
Ambassidae
Mugilidae
Polycentridae
Atheriniformes
Cichlidae-Pholidichthys
0.03 substitutions per site
%BS = 0-92
%BS = 93-99
90% Complete Matrix
ASTRAL-III Multispecies Coalescent
Atheriniformes
Blennioidei-Gramma
Embiocitidae
Pomacentridae
Ambassidae
Mugilidae
Congrogadidae
Pseudochromis
Plesiopidae
Polycentridae
Cichlidae-Pholidichthys
2.0 coalescent units
IQ-TREE Concatenated (Single Partition)
Plesiopidae
Congrogadidae
Pseudochromis
Pomacentridae
Embiocitidae
Blennioidei-Gramma
Ambassidae
Mugilidae
Polycentridae
Atheriniformes
Cichlidae-Pholidichthys
0.03 substitutions per site
IQ-TREE Concatenated (Multipartition)
Ambassidae
Mugilidae
Plesiopidae
Congrogadidae
Pseudochromis
Pomacentridae
Embiocitidae
Blennioidei-Gramma
Polycentridae
Atheriniformes
Cichlidae-Pholidichthys
0.03 substitutions per site
B
Pseudochromidae 100 100 100 100 100 100 30.2 62 31 62.3
Pomacentridae 100 100 100 100 100 100 32.2 76 31.4 75.8
Polycentridae 100 100 100 100 100 100 81.8 88 82.2 88.3
Plesiopidae 100 100 100 100 100 100 13.4 54.5 15 54.1
Mugilidae 100 100 100 100 100 100 68.8 83.4 68.7 84
Embiocitidae 100 100 100 100 100 100 80.9 94.9 81.4 94.8
Congrogadidae 100 100 100 100 100 100 63.7 86.8 63 87.7
Cichlidae-Pholidichthys 100 100 100 100 100 100 19 45 15.3 45.2
Blennioidei-Gramma 100 100 100 100 100 100 3.05 46 3.48 46.5
Atheriniformes 100 100 100 100 100 100 15.8 57.5 16.1 57.2
Ambassidae 100 100 100 100 100 100 68.6 81.6 66.9 81.5
ASTRAL Coalescent
Single Partition Ultrafast Bootstrap
Multiple Partition Ultrafast Bootstrap
ASTRAL Coalescent
Single Partition Ultrafast Bootstrap
Multiple Partition Ultrafast Bootstrap
gCF
sCF
gCF
sCF
75% Complete Matrix
90% Complete Matrix
75% Complete Matrix
90% Complete Matrix

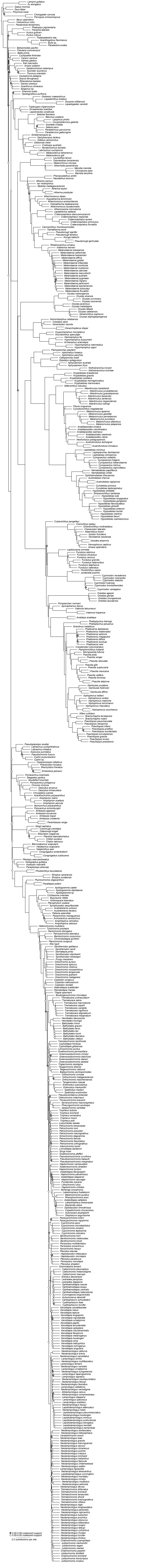

0.92-0.99 coalescent support
0.00-0.92 coalescent support
2.0 substitutions per site

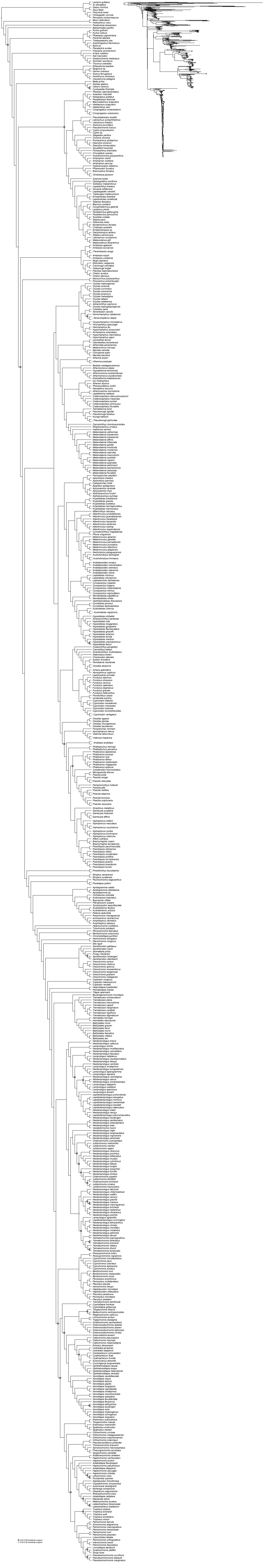
0.92-0.99 bootstrap support
0.00-0.92 bootstrap support

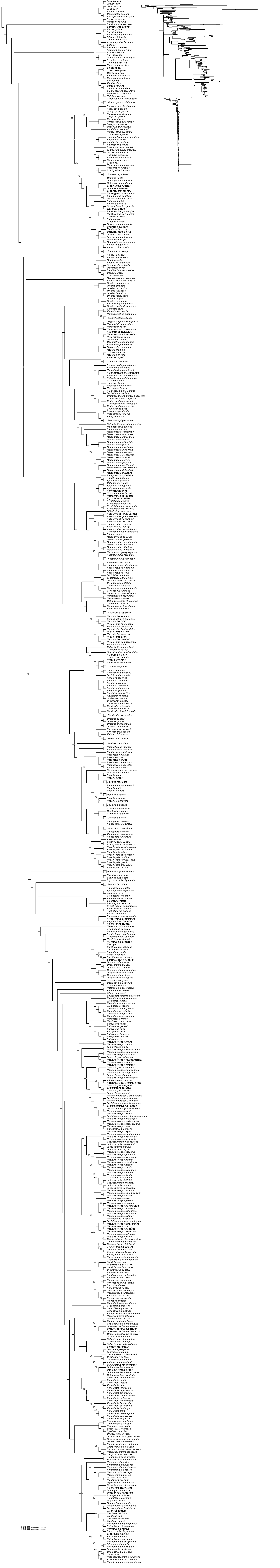

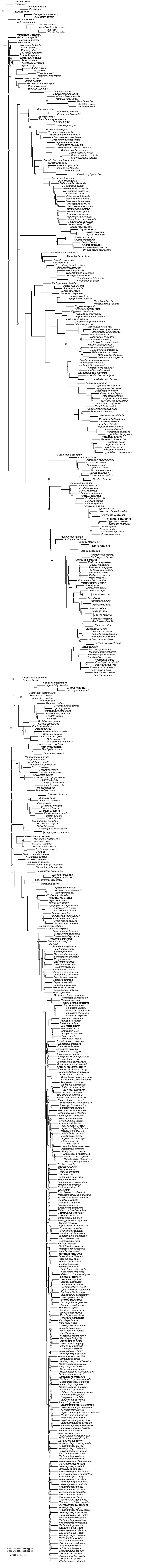
0.92-0.99 coalescent support
0.00-0.92 coalescent support
2.0 coalescent units

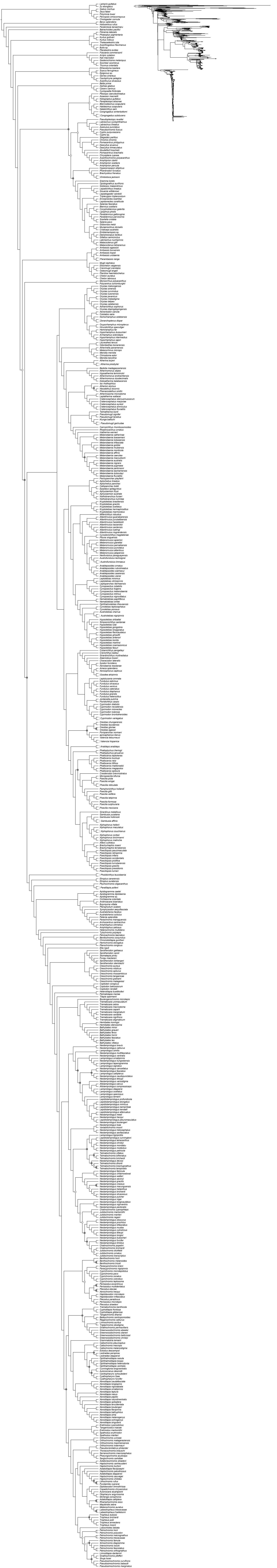

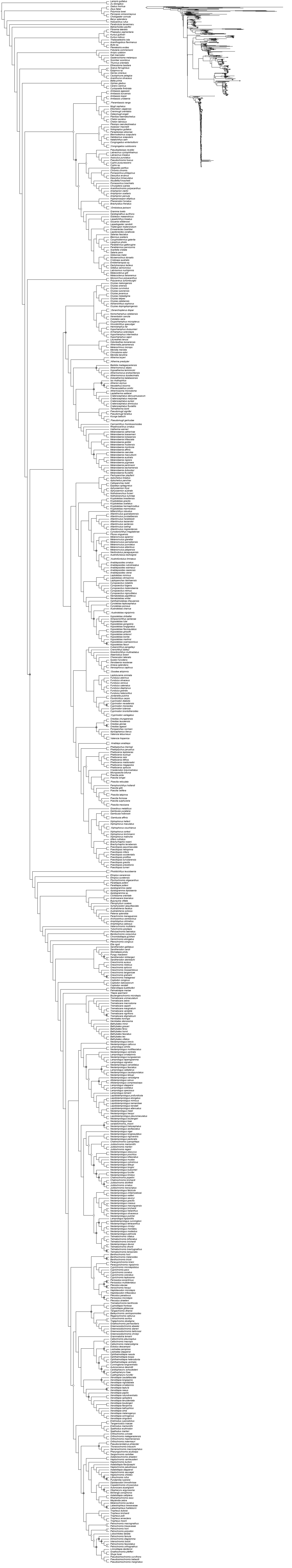

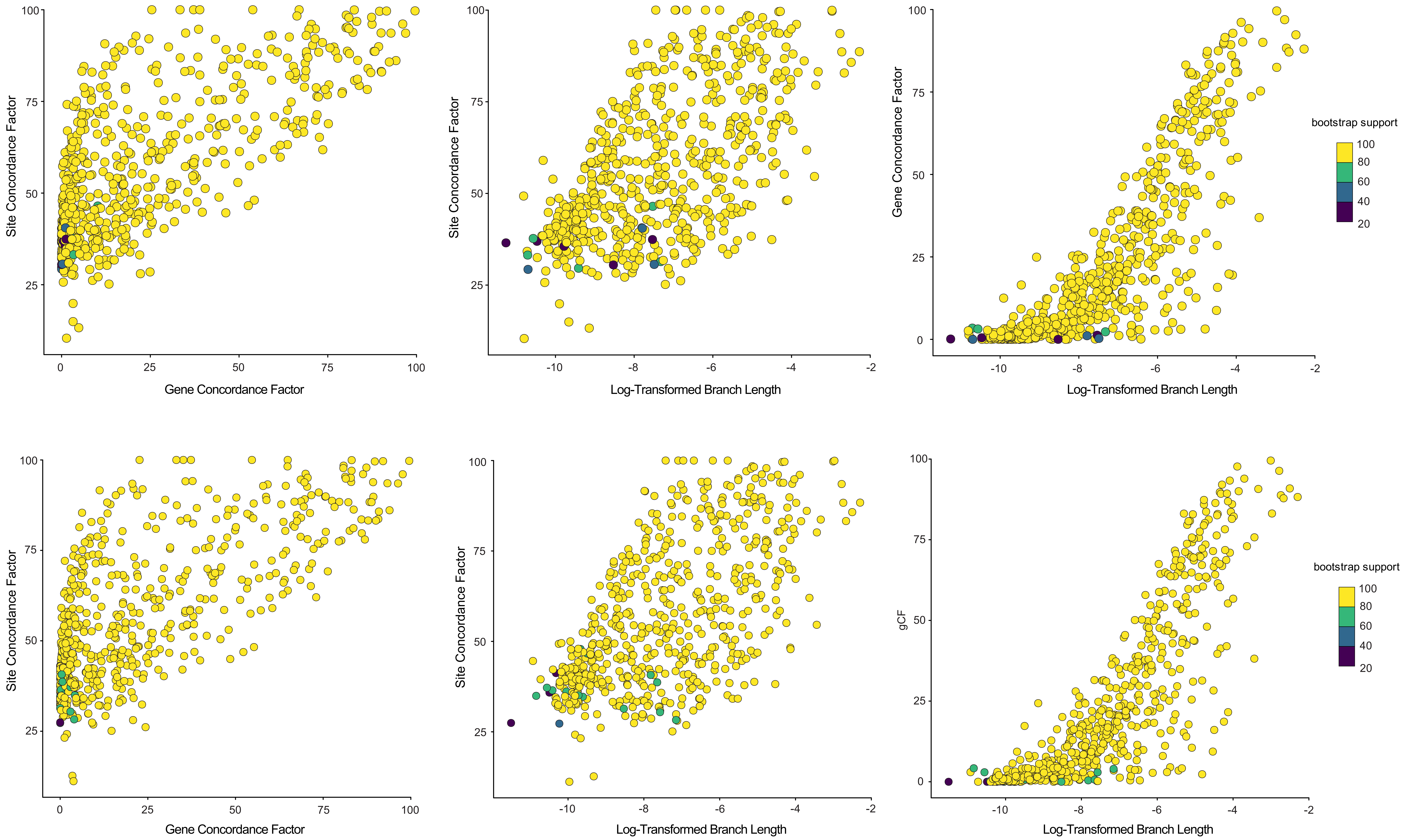
Site Concordance Factor
Gene Concordance Factor
Site Concordance Factor
Log-Transformed Branch Length
Gene Concordance Factor
Log-Transformed Branch Length
bootstrap support
100
80
60
40
20
Site Concordance Factor
Gene Concordance Factor
Site Concordance Factor
Log-Transformed Branch Length
gCF
Log-Transformed Branch Length
bootstrap support
100
80
60
40
20

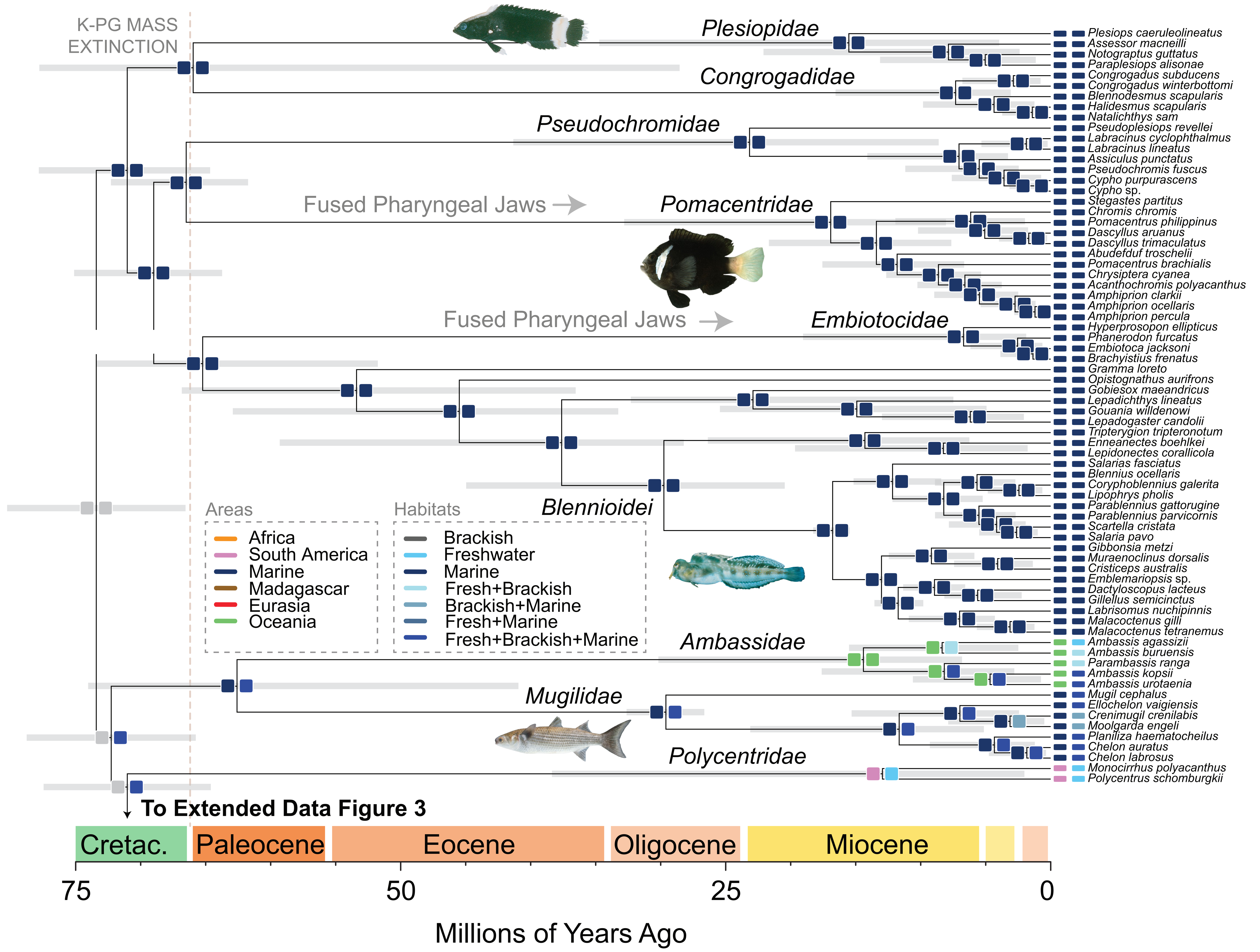

K-PG MASS EXTINCTION
Plesiopidae
Congrogadidae
Pseudochromidae
Fused Pharyngeal Jaws
Pomacentridae
Fused Pharyngeal Jaws
Embiotocidae
Blennioidei
Ambassidae
Mugilidae
Polycentridae
Areas
Africa
South America
Marine
Madagascar
Eurasia
Oceania
Habitats
Brackish
Freshwater
Marine
Fresh+Brackish
Brackish+Marine
Fresh+Marine
Fresh+Brackish+Marine
Plesiops caeruleolineatus
Assessor macneilli
Notograptus guttatus
Paraplesiops alisonae
Congrogadus subducens
Congrogadus winterbottomi
Blennodesmus scapularis
Halidesmus scapularis
Natalichthys sam
Pseudoplesiops revellei
Labracinus cyclophthalmus
Labracinus lineatus
Assiculus punctatus
Pseudochromis fuscus
Cypho purpurascens
Cypho sp.
Stegastes partitus
Chromis chromis
Pomacentrus philippinus
Dascyllus aruanus
Dascyllus trimaculatus
Abudefduf troschelii
Pomacentrus brachialis
Chrysiptera cyanea
Acanthochromis polyacanthus
Amphiprion clarkii
Amphiprion ocellaris
Amphiprion percula
Hyperprosopon ellipticus
Phanerodon furcatus
Embiotoca jacksoni
Brachyistius frenatus
Gramma loreto
Opistognathus aurifrons
Gobiesox maeandricus
Lepadichthys lineatus
Gouania willdenowi
Lepadogaster candolii
Tripterygion tripteronotum
Enneanectes boehlkei
Lepidonectes corallicola
Salarias fasciatus
Blennius ocellaris
Coryphoblennius galerita
Lipophrys pholis
Parablennius gattorugine
Parablennius parvicornis
Scartella cristata
Salaria pavo
Gibbonsia metzi
Muraenoclinus dorsalis
Cristiceps australis
Emblemariopsis sp.
Dactyloscopus lacteus
Gillellus semicinctus
Labrisomus nuchipinnis
Malacoctenus gilli
Malacoctenus tetranemus
Ambassis agassizii
Ambassis buruensis
Parambassis ranga
Ambassis kopsii
Ambassis urotaenia
Mugil cephalus
Ellochelon vaigiensis
Crenimugil crenilabis
Moolgarda engeli
Planiliza haematocheilus
Chelon auratus
Chelon labrosus
Monocirrhus polyacanthus
Polycentrus schomburgkii
To Extended Data Figure 3
Cretac.
Paleocene
Eocene
Oligocene
Miocene
75
50
25
0
Millions of Years Ago

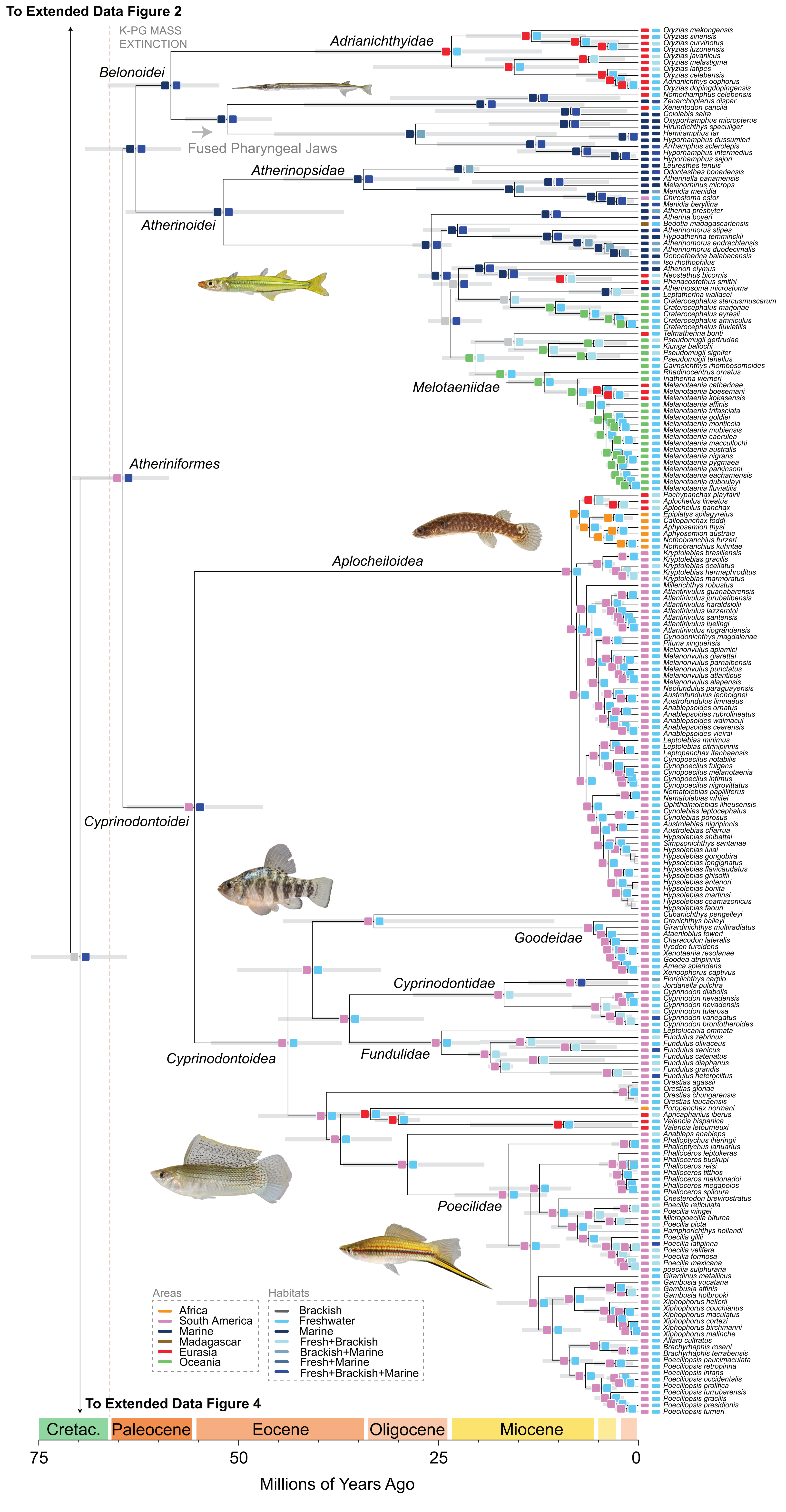

To Extended Data Figure 2
K-PG MASS EXTINCTION
Adrianichthyidae
Belonoidei
Fused Pharyngeal Jaws
Atherinopsidae
Atherinoidei
Melotaeniidae
Atheriniformes
Aplocheiloidea
Cyprinodontoidei
Goodeidae
Cyprinodontidae
Cyprinodontoidea
Fundulidae
Poecilidae
Oryzias mekongensis
Oryzias sinensis
Oryzias curvinotus
Oryzias luzonensis
Oryzias javanicus
Oryzias melastigma
Oryzias latipes
Oryzias celebensis
Adrianichthys oophorus
Oryzias dopingdopingensis
Nomorhamphus celebensis
Zenarchopterus dispar
Xenentodon cancila
Cololabis saira
Oxyporhamphus micropterus
Hirundichthys speculiger
Hemiramphus far
Hyporhamphus dussumieri
Arrhamphus sclerolepis
Hyporhamphus intermedius
Hyporhamphus sajori
Leuresthes tenuis
Odontesthes bonariensis
Atherinella panamensis
Melanorhinus microps
Menidia menidia
Chirostoma estor
Menidia beryllina
Atherina presbyter
Atherina boyeri
Bedotia madagascariensis
Atherinomorus stipes
Hypoatherina temminckii
Atherinomorus endrachtensis
Atherinomorus duodecimalis
Doboatherina balabacensis
Iso rhothophilus
Atherion elymus
Neostethus bicornis
Phenacostethus smithi
Atherinosoma microstoma
Leptatherina wallacei
Craterocephalus stercusmuscarum
Craterocephalus marjoriae
Craterocephalus eyresii
Craterocephalus amniculus
Craterocephalus fluviatilis
Telmatherina bonti
Pseudomugil gertrudae
Kiunga ballochi
Pseudomugil signifer
Pseudomugil tenellus
Cairnsichthys rhombosomoides
Rhadinocentrus ornatus
Iriatherina werneri
Melanotaenia catherinae
Melanotaenia boesemani
Melanotaenia kokasensis
Melanotaenia affinis
Melanotaenia trifasciata
Melanotaenia goldiei
Melanotaenia monticola
Melanotaenia mubiensis
Melanotaenia caerulea
Melanotaenia maccullochi
Melanotaenia australis
Melanotaenia nigrans
Melanotaenia pygmaea
Melanotaenia parkinsoni
Melanotaenia eachamensis
Melanotaenia duboulayi
Melanotaenia fluviatilis
Pachypanchax playfairii
Aplocheilus lineatus
Aplocheilus panchax
Epiplatys spilagyreius
Callopanchax toddi
Aphyosemion thysi
Aphyosemion australe
Nothobranchius furzeri
Nothobranchius kuhntae
Kryptolebias brasiliensis
Kryptolebias gracilis
Kryptolebias ocellatus
Kryptolebias hermaphroditus
Kryptolebias marmoratus
Millerichthys robustus
Atlantirivulus guanabarensis
Atlantirivulus jurubatibensis
Atlantirivulus haraldsiolii
Atlantirivulus lazzarotoi
Atlantirivulus santensis
Atlantirivulus luelingi
Atlantirivulus riograndensis
Cynodonichthys magdalenae
Pituna xinguensis
Melanorivulus apiamici
Melanorivulus giarettai
Melanorivulus parnaibensis
Melanorivulus punctatus
Melanorivulus atlanticus
Melanorivulus alapensis
Neofundulus paraguayensis
Austrofundulus leohoignei
Austrofundulus limnaeus
Anablepsoides ornatus
Anablepsoides rubrolineatus
Anablepsoides waimacui
Anablepsoides cearensis
Anablepsoides vieirai
Leptolebias minimus
Leptolebias citrinipinnis
Leptopanchax itanhaensis
Cynopoecilus notabilis
Cynopoecilus fulgens
Cynopoecilus melanotaenia
Cynopoecilus intimus
Cynopoecilus nigrovittatus
Nematolebias papilliferus
Nematolebias whitei
Ophthalmolebias ilheusensis
Cynolebias leptocephalus
Cynolebias porosus
Austrolebias nigripinnis
Austrolebias charrua
Hypsolebias shibattai
Simpsonichthys santanae
Hypsolebias lulai
Hypsolebias gongobira
Hypsolebias longignatus
Hypsolebias flavicaudatus
Hypsolebias ghisolfii
Hypsolebias antenori
Hypsolebias bonita
Hypsolebias martinsi
Hypsolebias coamazonicus
Hypsolebias faouri
Cubanichthys pengelleyi
Crenichthys baileyi
Girardinichthys multiradiatus
Ataeniobius toweri
Characodon lateralis
Ilyodon furcidens
Xenotaenia resolanae
Goodea atripinnis
Ameca splendens
Xenoophorus captivus
Floridichthys carpio
Jordanella pulchra
Cyprinodon diabolis
Cyprinodon nevadensis
Cyprinodon nevadensis
Cyprinodon tularosa
Cyprinodon variegatus
Cyprinodon brontotheroides
Leptolucania ommata
Fundulus zebrinus
Fundulus olivaceus
Fundulus xenicus
Fundulus catenatus
Fundulus diaphanus
Fundulus grandis
Fundulus heteroclitus
Orestias agassii
Orestias gloriae
Orestias chungarensis
Orestias laucaensis
Poropanchax normani
Apricaphanius iberus
Valencia hispanica
Valencia letourneuxi
Anableps anableps
Phalloptychus iheringii
Phalloptychus januarius
Phalloceros leptokeras
Phalloceros buckupi
Phalloceros reisi
Phalloceros titthos
Phalloceros maldonadoi
Phalloceros megapolos
Phalloceros spiloura
Cnesterodon brevirostratus
Poecilia reticulata
Poecilia wingei
Micropoecilia bifurca
Poecilia picta
Pamphorichthys hollandi
Poecilia gillii
Poecilia latipinna
Poecilia velifera
Poecilia formosa
Poecilia mexicana
poecilia sulphuraria
Girardinus metallicus
Gambusia yucatana
Gambusia affinis
Gambusia holbrooki
Xiphophorus hellerii
Xiphophorus couchianus
Xiphophorus maculatus
Xiphophorus cortezi
Xiphophorus birchmanni
Xiphophorus malinche
Alfaro cultratus
Brachyrhaphis roseni
Brachyrhaphis terrabensis
Poeciliopsis paucimaculata
Poeciliopsis retropinna
Poeciliopsis infans
Poeciliopsis occidentalis
Poeciliopsis prolifica
Poeciliopsis turrubarensis
Poeciliopsis gracilis
Poeciliopsis presidionis
Poeciliopsis turneri
Areas
Africa
South America
Marine
Madagascar
Eurasia
Oceania
Habitats
Brackish
Freshwater
Marine
Fresh+Brackish
Brackish+Marine
Fresh+Marine
Fresh+Brackish+Marine
To Extended Data Figure 4
Cretac.
Paleocene
Eocene
Oligocene
Miocene
75
50
25
0
Millions of Years Ago

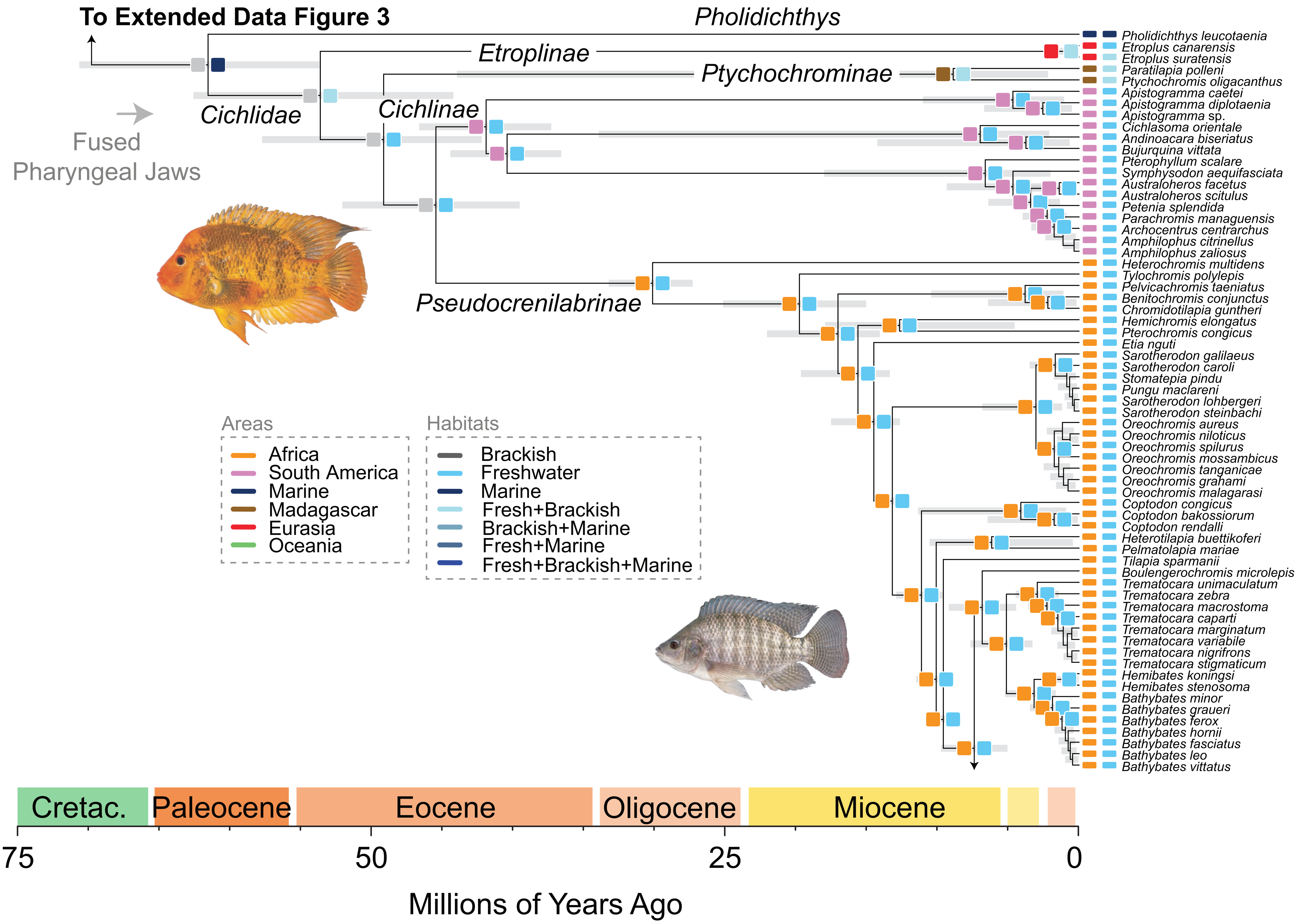

To Extended Data Figure 3
Pholidichthys
Etroplinae
Ptychochrominae
Cichlidae
Cichlinae
Fused
Pharyngeal Jaws
Pseudocrenilabrinae
Areas
Africa
South America
Marine
Madagascar
Eurasia
Oceania
Habitats
Brackish
Freshwater
Marine
Fresh+Brackish
Brackish+Marine
Fresh+Marine
Fresh+Brackish+Marine
Pholidichthys leucotaenia
Etroplus canarensis
Etroplus suratensis
Paratilapia polleni
Ptychochromis oligacanthus
Apistogramma caetei
Apistogramma diplotaenia
Apistogramma sp.
Cichlasoma orientale
Andinoacara biseriatus
Bujurquina vittata
Pterophyllum scalare
Symphysodon aequifasciata
Australoheros facetus
Australoheros scitulus
Petenia splendida
Parachromis managuensis
Archocentrus centrarchus
Amphilophus citrinellus
Amphilophus zaliosus
Heterochromis multidens
Tylochromis polylepis
Pelvicachromis taeniatus
Benitochromis conjunctus
Chromidotilapia guntheri
Hemichromis elongatus
Pterochromis congicus
Etia nguti
Sarotherodon galilaeus
Sarotherodon caroli
Stomatepia pindu
Pungu maclareni
Sarotherodon lohbergeri
Sarotherodon steinbachi
Oreochromis aureus
Oreochromis niloticus
Oreochromis spilurus
Oreochromis mossambicus
Oreochromis tanganicae
Oreochromis grahami
Oreochromis malagarasi
Coptodon congicus
Coptodon bakossiorum
Coptodon rendalli
Heterotilapia buettikoferi
Pelmatolapia mariae
Tilapia sparmanii
Boulengerochromis microlepis
Trematocara unimaculatum
Trematocara zebra
Trematocara macrostoma
Trematocara caparti
Trematocara marginatum
Trematocara variabile
Trematocara nigrifrons
Trematocara stigmaticum
Hemibates koningsi
Hemibates stenosoma
Bathybates minor
Bathybates graueri
Bathybates ferox
Bathybates hornii
Bathybates fasciatus
Bathybates leo
Bathybates vittatus
Cretac.
Paleocene
Eocene
Oligocene
Miocene
75
50
25
0
Millions of Years Ago

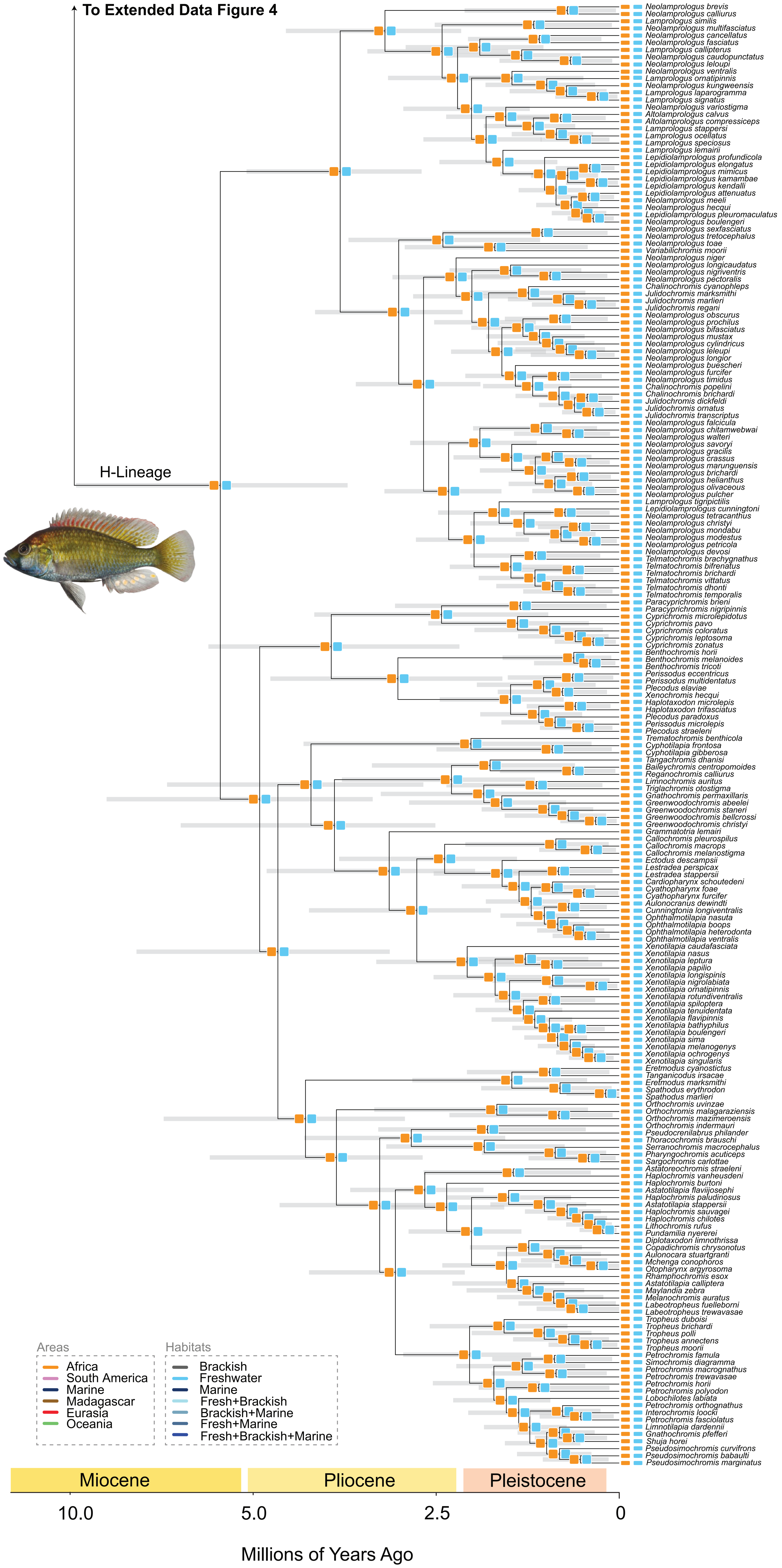

To Extended Data Figure 4
H-Lineage
Neolamprologus brevis
Neolamprologus calliurus
Lamprologus similis
Neolamprologus multifasciatus
Neolamprologus cancellatus
Neolamprologus fasciatus
Lamprologus callipterus
Neolamprologus caudopunctatus
Neolamprologus leloupi
Neolamprologus ventralis
Lamprologus ornatipinnis
Neolamprologus kungweensis
Lamprologus laparogramma
Lamprologus signatus
Neolamprologus variostigma
Altolamprologus calvus
Altolamprologus compressiceps
Lamprologus stappersi
Lamprologus ocellatus
Lamprologus speciosus
Lamprologus lemairii
Lepidiolamprologus profundicola
Lepidiolamprologus elongatus
Lepidiolamprologus mimicus
Lepidiolamprologus kamambae
Lepidiolamprologus kendalli
Lepidiolamprologus attenuatus
Neolamprologus meeli
Neolamprologus hecqui
Lepidiolamprologus pleuromaculatus
Neolamprologus boulengeri
Neolamprologus sexfasciatus
Neolamprologus tretocephalus
Neolamprologus toae
Variabilichromis moorii
Neolamprologus niger
Neolamprologus longicaudatus
Neolamprologus nigriventris
Neolamprologus pectoralis
Chalinochromis cyanophleps
Julidochromis marksmithi
Julidochromis marlieri
Julidochromis regani
Neolamprologus obscurus
Neolamprologus prochilus
Neolamprologus bifasciatus
Neolamprologus mustax
Neolamprologus cylindricus
Neolamprologus leleupi
Neolamprologus longior
Neolamprologus buescheri
Neolamprologus furcifer
Neolamprologus timidus
Chalinochromis popelini
Chalinochromis brichardi
Julidochromis dickfeldi
Julidochromis ornatus
Julidochromis transcriptus
Neolamprologus falcicula
Neolamprologus chitamwebwai
Neolamprologus walteri
Neolamprologus savoryi
Neolamprologus gracilis
Neolamprologus crassus
Neolamprologus marunguensis
Neolamprologus brichardi
Neolamprologus helianthus
Neolamprologus olivaceous
Neolamprologus pulcher
Lamprologus tigripictilis
Lepidiolamprologus cunningtoni
Neolamprologus tetracanthus
Neolamprologus christyi
Neolamprologus mondabu
Neolamprologus modestus
Neolamprologus petricola
Neolamprologus devosi
Telmatochromis brachygnathus
Telmatochromis bifrenatus
Telmatochromis brichardi
Telmatochromis vittatus
Telmatochromis dhonti
Telmatochromis temporalis
Paracyprichromis brieni
Paracyprichromis nigripinnis
Cyprichromis microlepidotus
Cyprichromis pavo
Cyprichromis coloratus
Cyprichromis leptosoma
Cyprichromis zonatus
Benthochromis horii
Benthochromis melanoides
Benthochromis tricoti
Perissodus eccentricus
Perissodus multidentatus
Plecodus elaviae
Xenochromis hecqui
Haplotaxodon microlepis
Haplotaxodon trifasciatus
Plecodus paradoxus
Perissodus microlepis
Plecodus straeleni
Trematochromis benthicola
Cyphotilapia frontosa
Cyphotilapia gibberosa
Tangachromis dhanisi
Baileychromis centropomoides
Reganochromis calliurus
Limnochromis auritus
Triglachromis otostigma
Gnathochromis permaxillaris
Greenwoodochromis abeelei
Greenwoodochromis staneri
Greenwoodochromis bellcrossi
Greenwoodochromis christyi
Grammatotria lemairi
Callochromis pleurospilus
Callochromis macrops
Callochromis melanostigma
Ectodus descampsii
Lestradea perspicax
Lestradea stappersii
Cardiopharynx schoutedeni
Cyathopharynx foae
Cyathopharynx furcifer
Aulonocranus dewindti
Cunningtonia longiventralis
Ophthalmotilapia nasuta
Ophthalmotilapia boops
Ophthalmotilapia heterodonta
Ophthalmotilapia ventralis
Xenotilapia caudafasciata
Xenotilapia nasus
Xenotilapia leptura
Xenotilapia papilio
Xenotilapia longispinis
Xenotilapia nigrolabiata
Xenotilapia ornatipinnis
Xenotilapia rotundiventralis
Xenotilapia spiloptera
Xenotilapia tenuidentata
Xenotilapia flavipinnis
Xenotilapia bathyphilus
Xenotilapia boulengeri
Xenotilapia sima
Xenotilapia melanogenys
Xenotilapia ochrogenys
Xenotilapia singularis
Eretmodus cyanostictus
Tanganicodus irsacae
Eretmodus marksmithi
Spathodus erythrodon
Spathodus marlieri
Orthochromis uvinzae
Orthochromis malagaraziensis
Orthochromis mazimeroensis
Orthochromis indermauri
Pseudocrenilabrus philander
Thoracochromis brauschi
Serranochromis macrocephalus
Pharyngochromis acuticeps
Sargochromis carlottae
Astatoreochromis straeleni
Haplochromis vanheusdeni
Haplochromis burtoni
Astatotilapia flaviijosephi
Haplochromis paludinosus
Astatotilapia stappersii
Haplochromis sauvagei
Haplochromis chilotes
Lithochromis rufus
Pundamilia nyererei
Diplotaxodon limnothrissa
Copadichromis chrysonotus
Aulonocara stuartgranti
Mchenga conophoros
Otopharynx argyrosoma
Rhamphochromis esox
Astatotilapia calliptera
Maylandia zebra
Melanochromis auratus
Labeotropheus fuelleborni
Labeotropheus trewavasae
Tropheus duboisi
Tropheus brichardi
Tropheus polli
Tropheus annectens
Tropheus moorii
Petrochromis famula
Simochromis diagramma
Petrochromis macrognathus
Petrochromis trewavasae
Petrochromis horii
Petrochromis polyodon
Lobochilotes labiata
Petrochromis orthognathus
Interochromis loocki
Petrochromis fasciolatus
Limnotilapia dardennii
Gnathochromis pfefferi
Shuja horei
Pseudosimochromis curvifrons
Pseudosimochromis babaulti
Pseudosimochromis marginatus
Areas
Africa
South America
Marine
Madagascar
Eurasia
Oceania
Habitats
Brackish
Freshwater
Marine
Fresh+Brackish
Brackish+Marine
Fresh+Marine
Fresh+Brackish+Marine
Miocene
Pliocene
Pleistocene
10.0
5.0
2.5
0
Millions of Years Ago

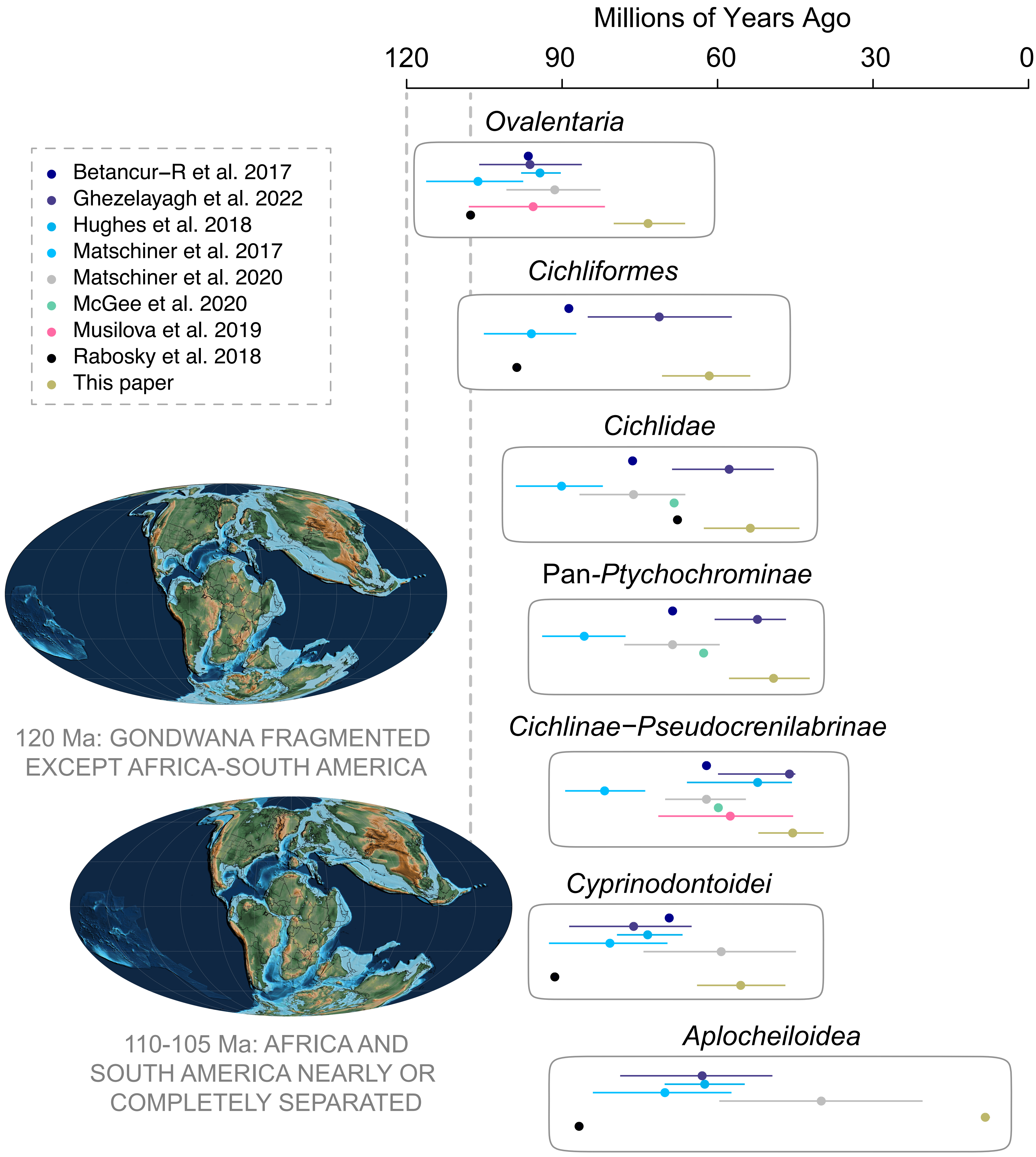

Millions of Years Ago
120
90
60
30
0
Ovalentaria
Cichliformes
Cichlidae
Pan-Ptychochrominae
Cichlinae−Pseudocrenilabrinae
Cyprinodontoidei
Aplocheiloidea
Betancur−R et al. 2017
Ghezelayagh et al. 2022
Hughes et al. 2018
Matschiner et al. 2017
Matschiner et al. 2020
McGee et al. 2020
Musilova et al. 2019
Rabosky et al. 2018
This paper
120 Ma: GONDWANA FRAGMENTED
EXCEPT AFRICA-SOUTH AMERICA
110-105 Ma: AFRICA AND
SOUTH AMERICA NEARLY OR
COMPLETELY SEPARATED

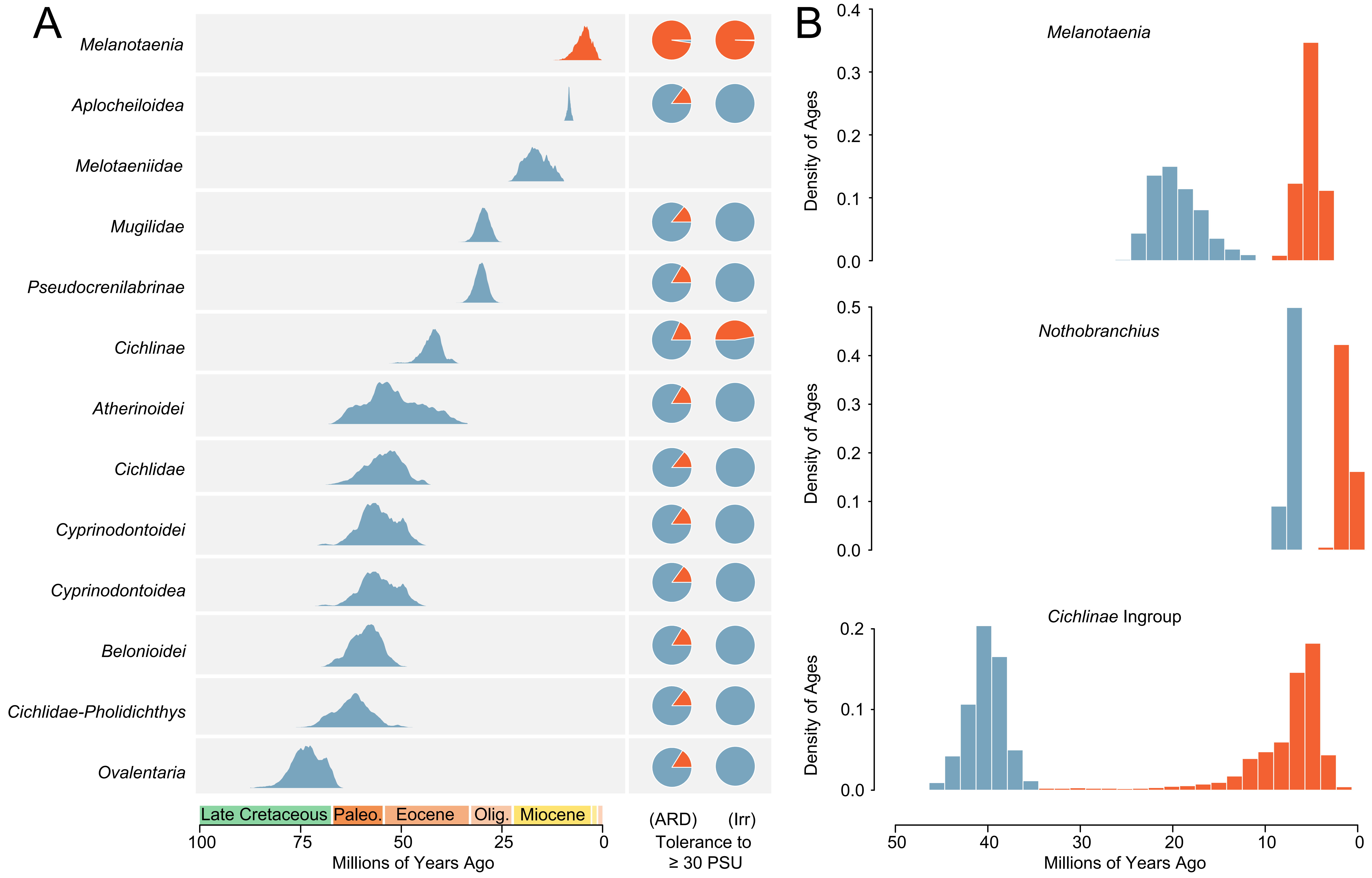
A
Melanotaenia
Aplocheiloidea
Melotaeniidae
Mugilidae
Pseudocrenilabrinae
Cichlinae
Atherinoidei
Cichlidae
Cyprinodontoidei
Cyprinodontoidea
Belonioidei
Cichlidae-Pholidichthys
Ovalentaria
Late Cretaceous
Paleo.
Eocene
Olig.
Miocene
100
75
50
25
0
Millions of Years Ago
(ARD)
(Irr)
Tolerance to
≥ 30 PSU
B
Melanotaenia
Nothobranchius
Cichlinae Ingroup
Density of Ages
0.4
0.3
0.2
0.1
0.0
0.5
50
40
30
20
10
0
Millions of Years Ago

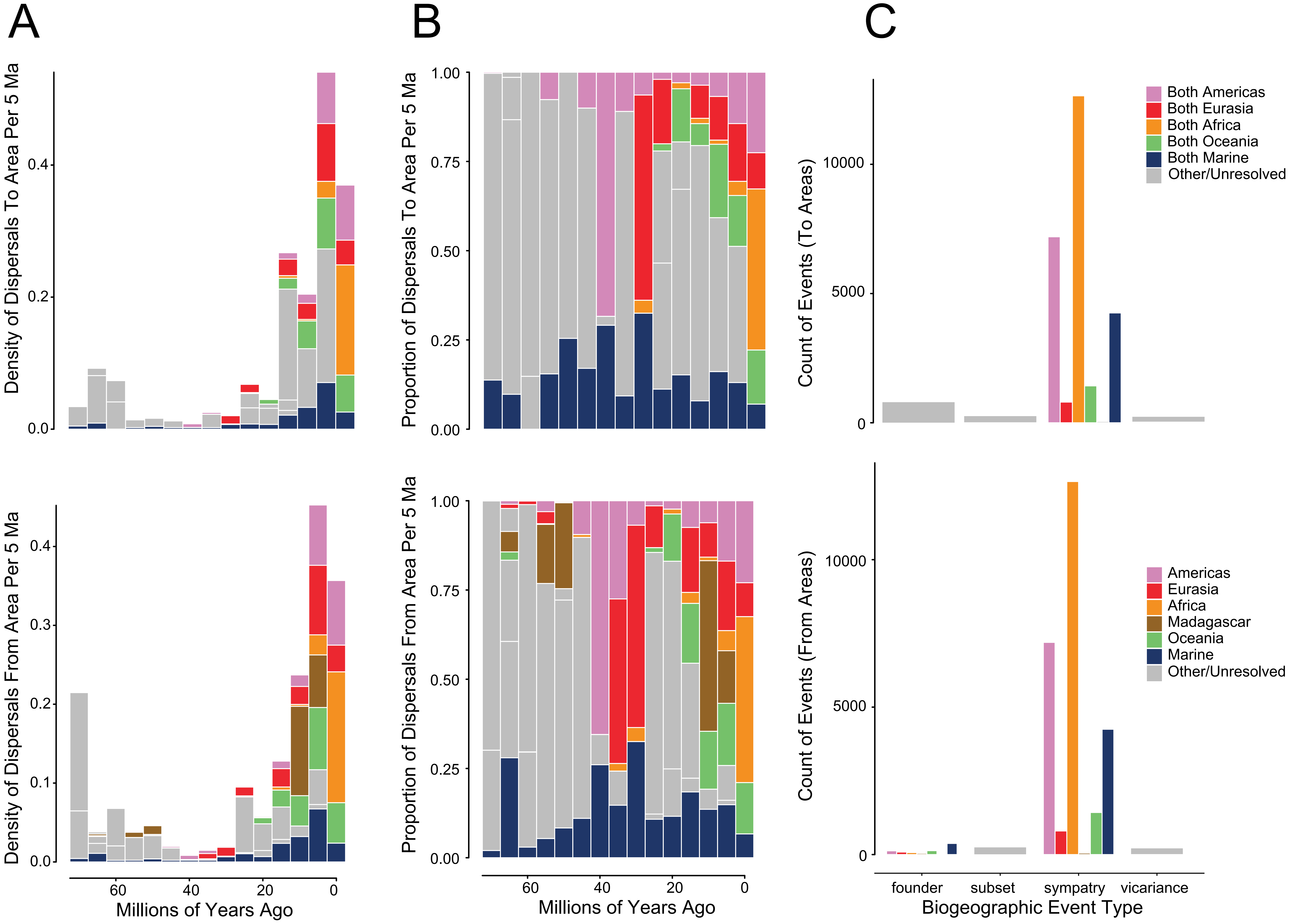
A
B
C
Density of Dispersals To Area Per 5 Ma
Density of Dispersals From Area Per 5 Ma
Proportion of Dispersals To Area Per 5 Ma
Proportion of Dispersals From Area Per 5 Ma
Count of Events (To Areas)
Count of Events (From Areas)
Millions of Years Ago
Biogeographic Event Type
founder
subset
sympatry
vicariance
Both Americas
Both Eurasia
Both Africa
Both Oceania
Both Marine
Other/Unresolved
Americas
Eurasia
Africa
Madagascar
Oceania
Marine
Other/Unresolved

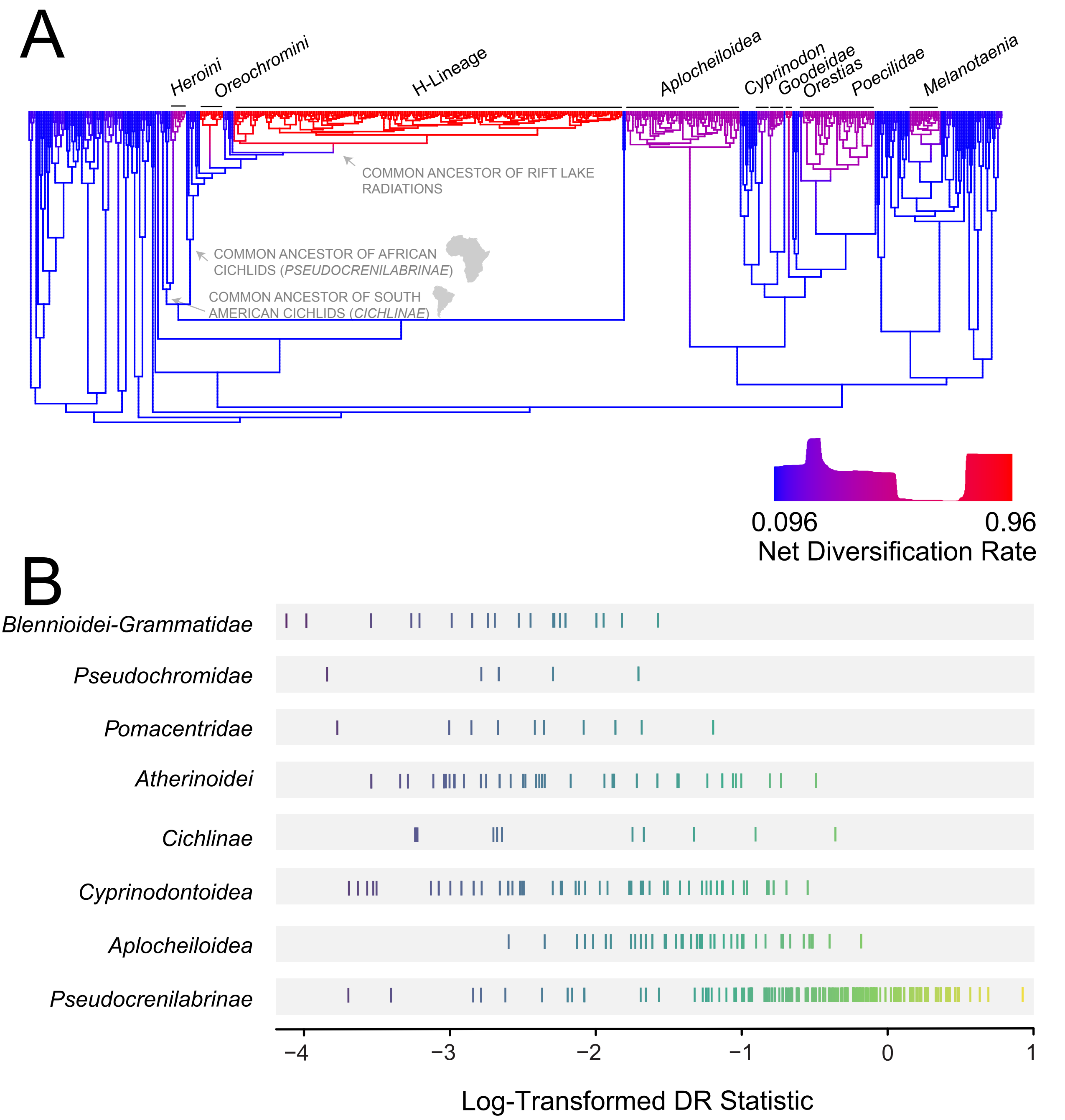
A
Heroini
Oreochromini
H-Lineage
Aplocheiloidea
Cyprinodon
Goodeidae
Orestias
Poecilidae
Melanotaenia
COMMON ANCESTOR OF RIFT LAKE RADIATIONS
COMMON ANCESTOR OF AFRICAN CICHLIDS (PSEUDOCRENILABRINAE)
COMMON ANCESTOR OF SOUTH AMERICAN CICHLIDS (CICHLINAE)
0.096
0.96
Net Diversification Rate
B
Blennioidei-Grammatidae
Pseudochromidae
Pomacentridae
Atherinoidei
Cichlinae
Cyprinodontoidea
Aplocheiloidea
Pseudocrenilabrinae
−4
−3
−2
−1
0
1
Log-Transformed DR Statistic

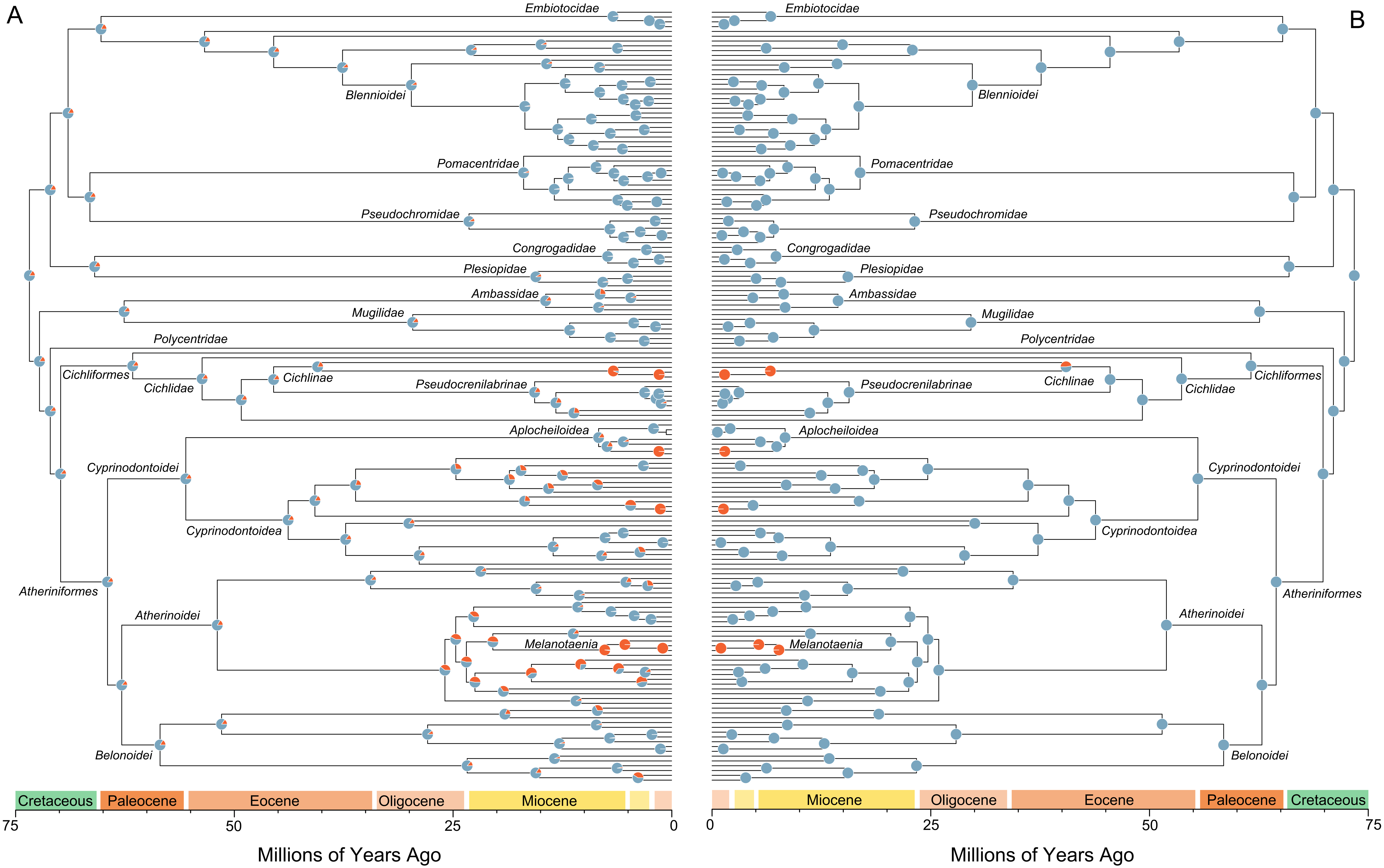
A
B
Embiotocidae
Blennioidei
Pomacentridae
Pseudochromidae
Congrogadidae
Plesiopidae
Ambassidae
Mugilidae
Polycentridae
Cichliformes
Cichlidae
Cichlinae
Pseudocrenilabrinae
Aplocheiloidea
Cyprinodontoidei
Cyprinodontoidea
Atheriniformes
Atherinoidei
Melanotaenia
Belonoidei
Cretaceous
Paleocene
Eocene
Oligocene
Miocene
75
50
25
0
Millions of Years Ago
